# A computational pipeline for spatial mechano-transcriptomics

**DOI:** 10.1101/2023.08.03.551894

**Authors:** Adrien Hallou, Ruiyang He, Benjamin D. Simons, Bianca Dumitrascu

## Abstract

Advances in spatial profiling technologies are providing insights into how molecular programs are influenced by local signaling and environmental cues. However, cell fate specification and tissue patterning involve the interplay of biochemical and mechanical feedback. Here, we develop a computational framework that enables the joint statistical analysis of transcriptional and mechanical signals in the context of spatial transcriptomics. To illustrate the application and utility of the approach, we use spatial transcriptomics data from the developing mouse embryo to infer the forces acting on individual cells, and use these results to identify mechanical, morphometric, and gene expression signatures that are predictive of tissue compartment boundaries. In addition, we use geoadditive structural equation modeling to identify gene modules that predict the mechanical behavior of cells in an unbiased manner. This computational framework is easily generalized to other spatial profiling contexts, providing a generic scheme for exploring the interplay of biomolecular and mechanical cues in tissues.

## Introduction

The advent of single-cell profiling technologies has transformed our understanding of the mechanisms that control cell state transitions and lineage hierarchies. Through the development of computational and statistical methods, these approaches provide a window into the transcriptional and epigenetic programs that control cell fate under normal and perturbed conditions. However, these programs do not act in isolation, but respond to environmental cues and collective cell behaviors, mediated by reciprocal signaling networks as well as mechanical forces and their coupling through mechanochemical feedback loops[1, 2, 3, 4]. In recent years, the advent of spatial -omics techniques [5] has enabled the profiling of gene expression [6, 7], protein composition [8] and chromatin accessibility [9] at single cell resolution in whole embryos and tissue sections opening a window on the correlation between cell state and spatial cues.

Despite their promise, spatial -omics methods and associated computational analysis pipelines currently struggle to integrate molecular profiling measures with interpretable cell morphology metrics and local mechanical forces. While high-throughput sequencing technologies such as Slide-seq [10] enable coverage of the entire transcriptome, they are limited to supra-cellular spatial resolution and as such, cannot recover such information. In contrast, ISH-based methods such as seqFISH [11, 12] and MERFISH [13, 14], offer a more limited transcriptomic coverage, yet provide single cell or even subcellular spatial resolution in addition to access to cell morphology. Indeed, immunostaining of transmembrane proteins allows segmentation of cell contours and extraction of whole-cell morphometric measures. Such cellular morphologies have been used either alone, for cell type classification and pseudotime inference [15], or in combination with gene expression data, for cell type clustering refinement [16, 17] and cross-modality prediction [18].

However, computational frameworks for linking genomics to tissue-level mechanical signatures such as tension at cell-cell junctions, strain and stress are currently lacking. Indeed, when considering the joint modeling of gene expression and morphology, existing approaches target only ’local’ morphological properties (e.g. cell roundness or volume), treating cells as independent from each other and from their spatial environment. Since mechanical morphometrics are global properties of cellular aggregates, new methods are needed to estimate these quantities from images and to test for associations between them and gene expression signatures in the presence of spatial confounders.

Here, to meet this challenge, we introduce a joint spatial mechano-transcriptomics framework to investigate simultaneously the transcriptional, morphological and mechanical state of cells in a tissue context at single-cell resolution. To develop this method, we make use of image-based mechanical force inference, an approach rooted in the physics of cellular materials [19]. To illustrate the potential of this approach, we use it to quantify tensions at cell-cell junctions and intracellular pressure in the context of multicellular tissues [20, 21, 22, 23]. In particular, we use selected regions of an embryonic day (E)8.5 mouse embryo spatially profiled using seqFISH [24]. We show that, by integrating transcriptomic profiling with local mechanical measures, we can gain insight into the mechanisms that promote boundary formation during development, as well as the role of mechano-responsive regulatory pathways in driving cell segregation and spatial patterning. We investigate the relationship between transcriptional profiles and mechanical forces at the single cell level, demonstrating the existence of gene modules whose expression patterns are significantly associated with the mechanical state of the cell, while accounting for spatial confounders. Finally, exploring higher order interactions between gene expression and mechanics, we show that mechano-associated genes display a variety of non-linear responses to mechanical signals. Overall, this study provides a computational framework to investigate mechanobiology in an unbiased manner, offering the potential to uncover the directional relationships between mechanical forces and gene expression in a spatial context, identify candidate mechanosensors or mechano-effectors, and delineate mechanical and mechanochemical feedback loops involved in cell fate decisions, pattern formation and tissue morphogenesis.

## Results

### An integrated mechano-transcriptomics analysis of mouse organogenesis

Can the gene expression signature of cells provide information on the local mechanical forces that act upon them? How does the interaction between genomics and mechanics inform the acquisition of cell identity and establishment of tissue compartments in developmental contexts? To begin to answer these questions, we developed a multi-step computational framework based on spatial transcriptomics for the integrated statistical analysis of mechanical forces and gene expression at cellular resolution. As shown in Figure 1A, our approach combines three main steps: data processing and image analysis; streamlined image-based mechanical force inference; and joint mechanical and transcriptomic statistical analysis.

**Figure 1:**
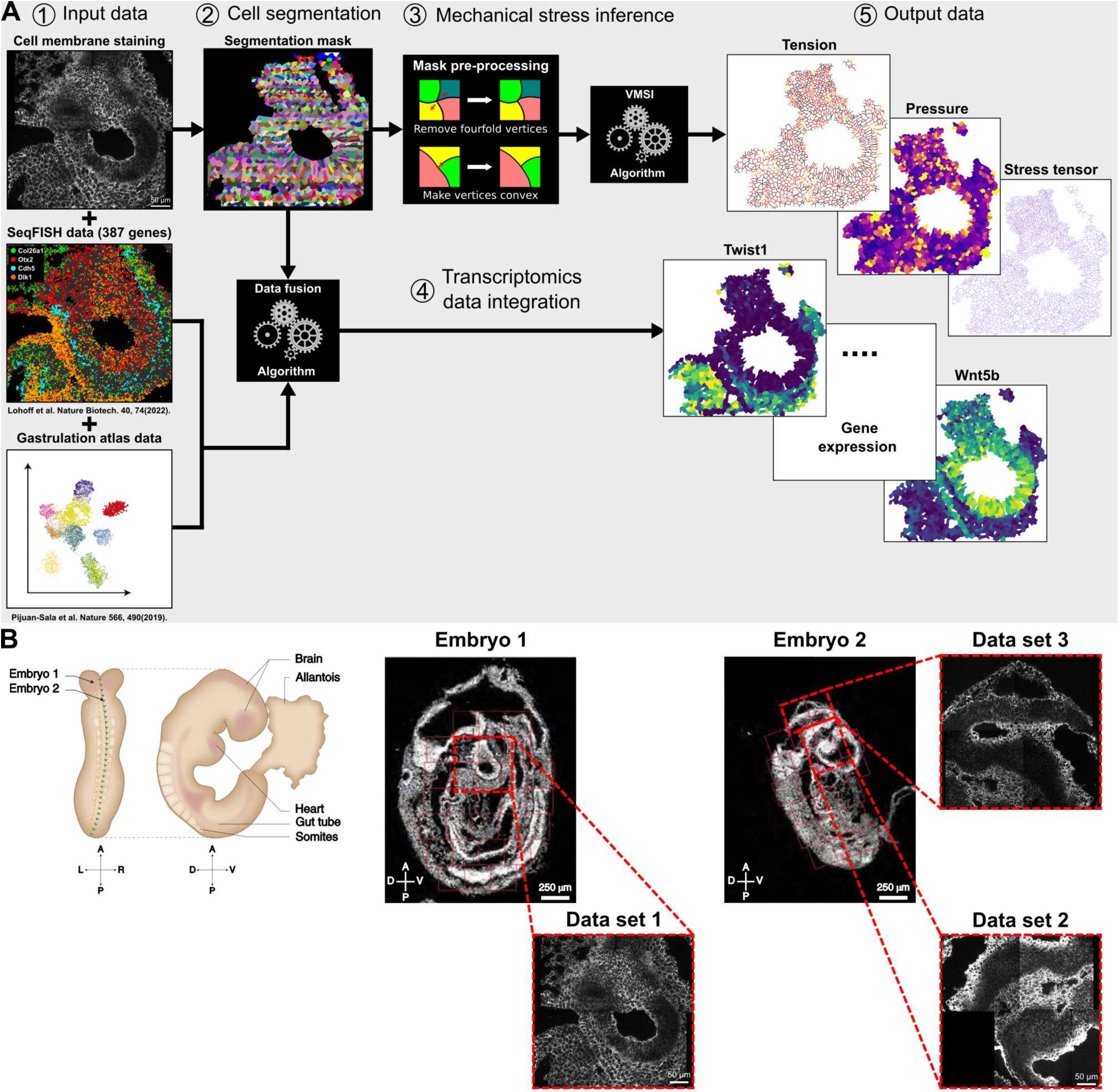
A spatial mechano-transcriptomics framework applied to a E8.5 mouse embryo seqFISH data set. **A:** Schematic describing our proposed framework for spatial mechano-transcriptomics data acquisition and processing. 1. Input data is composed of cell membrane immunostainings and seqFISH images of sagital sections of E8.5 mouse embryos and scRNA-Seq data from the mouse gastrulation atlas. 2. Instance segmentation masks of cell contours obtained using a deep learning segmentation pipeline. 3.Cell segmentation masks pre-processing and automated tension and pressure maps computation. 4.Generation of inputed gene expression levels for the whole mouse transcriptome for each segmented cell. 5.Output data of the framework composed of tension, pressure and mechanical stress tensor spatial maps and spatial maps of inputed gene expression for the whole murine transcriptome. **B**: Schematic (top left) and images of the two different embryo sagital sections considered in this study with close-ups of the 3 different brain regions studied in more details thereafter. Data set 1: Forebrain/Midbrain/Hindbrain (FMH) and Neural Crest (NC) regions of Embryo 1 brain; data set 2: Cranial Mesoderm (CM) and FHM regions of Embryo 2 brain; and data set 3: Midbrain/Hindbrain boundary (MHB) region of Embryo 2 brain.

As a starting point, we take as input images of tissue or embryo sections where cell membranes have been labelled with fluorescent markers (see e.g. the spatial transcriptomics seqFISH data set in Figure 1B). The fluorescent markers enable image-based segmentation of cell contours as well as the quantification and spatial localisation of selected transcripts at cellular resolution. Based on this analysis, we then generate segmentation masks with annotated coordinates of cell-cell junctions and vertices (see Methods). To infer the mechanical forces that act upon cells, we will take advantage of a mechanical force inference approach [23]. To safely apply this method, it is necessary to recover information on precise cellular shapes and physical cell-cell contacts, calling for high quality image segmentation. Moreover, to fulfil the constraints of our inference method, four-fold vertices – junctions that are shared between four neighbouring cells — must be reconciled and removed while cell edges must be convex at vertices (see Methods). The resulting spatial mask serves as input into an image-based mechanical force inference pipeline (Figure 1A and Figure Sup.1A).

Different algorithms exist for image-based force inference [23]. Here, we chose to implement the VMSI approach proposed by Noll *et al.* [22]. This algorithm uses a non-planar triangulation of junctional tensions to form a dual representation of the cell array geometry. A simultaneous fit of junctions with circular arcs then allows the inference of both tensions and cellular pressures up to a multiplicative and additive constant, respectively (see Methods). In doing so, it exhibits both increased accuracy and robustness compared to other force inference methods [23], particularly when the pressure differential between adjacent cells is large. We benchmark and calibrate this variational method for mechanical force inference against a variety of optimisers, choices of hyperparameters in real data and simulations (Figure Sup.1B-G), ensuring it is robust to perturbations and noise sources encountered in experimental data, and make it available as a python package. Here, we have expanded the utility of the original mechanical stress inference method by providing improved quality control tools for the resolution of “invalid” vertices and options for image tiling for large images (see Methods). The mechanical stress inference pipeline provides as output inferred intracellular pressures, tensions at cell-cell junctions and mechanical stress tensors for each segmented cell in the image. Both scalar and tensor quantities are determined. Scalar quantities are directly output as features, while tensorial quantities, such as the mechanical stress tensor, are converted to features that summarise the eigenvectors, orientation, and anisotropy of the tensor. This ensures that all resulting features are independently interpretable (see Methods).

Alongside the measured transcriptomic readouts, the mechanical estimates – tensions, pressures, stress tensor – comprise a mosaic representation of spatial cellular identity. We use these interpretable features to quantify statistical associations between genomic and mechanical measures. Using this approach, we can then build structural equation models which take into account spatial confounders [25], and identify known mechanosensors as well as genes and ligand-receptor (LR) pairs associated with cell-cell junctional tension variability along tissue compartment boundaries.

### Boundaries between tissue compartments are characterized by both gene expression and elevated interfacial tension

To illustrate the application and potential of this approach, we first apply our pipeline to the study of boundary formation in the gastrulating mouse embryo. The mechanisms that drive the formation of precise boundaries between tissue compartments in the developing embryo have been the subject of long-standing interest and debate [26, 27]. Does cell fate specification precede a phase of cell rearrangement and boundary formation, or does the positioning of cells induce cell fate acquisition? In the context of cell sorting, emphasis has been placed on the ability of cells to discriminate contacts between cells of the same cell-type – homotypic contacts – and between cells of a different cell-type – heterotypic contacts [28, 29]. Evidence for this phenomenon was first shown in pioneering work by Townes and Holtfreter, who demonstrated that mixed dissociated cells from different embryonic regions could progressively sort into segregated cell clusters [30].

Various hypotheses have been proposed to explain the basis of this phenomenon, either through differential cell adhesion (also known as the Differential Adhesion Hypothesis – DAH) [31], preferential cell adhesion (also known as Selective Adhesion hypothesis – SAH) [32], differential cell contractility (also known as Differential Interfacial Tension Hypothesis – DITH) [33] or juxtacrine signaling generating cell-cell repulsion at heterotypic cell contacts (also known as Higher Interfacial Tensions – HIT) [34, 35]. Based on modelling based approaches and experimental studies [36, 37, 38], it was established that both cell-cell adhesion and cell contractility contribute to the tuning of a single physical quantity, the cell-cell junctional tension (also known as interfacial tension or contact tension), which is the quantity that is directly inferred with our image-based force inference algorithm. Thus, it is possible to formulate the four hypotheses above in terms of cell-cell junctional tension. To understand how, consider two different cell-types, A and B, displaying homotypic junctional tensions between cells of the same type, *T_AA_* and *T_BB_* respectively, and an heterotypic junctional tension, *T_AB_*, between cells of different types. For the boundary to be maintained between segregated populations of A and B type cells, or for A and B cells to segregate if initially mixed, DAH and DITH require that *T_AA_ > T_AB_ > T_BB_*, whereas SAH and HIT require that *T_AB_ >* max(*T_AA_*, *T_BB_*). Therefore, using a combination of cell type annotation based on the transcriptomics data and the results of the mechanical force inference analysis, it should be possible to distinguish between these two scenarios.

To test our framework, we applied the force inference pipeline to three published spatial transcriptomics data sets of the embryonic day E8.5 mouse embryo obtained using the seqFISH pipeline [24] and shown in Figure 1A. We focused attention on three examples of boundary formation (Figure 3A), where we could distinguish between distinct cell types based on their transcriptional signature: Data set 1 shows a boundary between cells with a neural crest signature and the Forebrain/Midbrain/Hindbrain (FMH), two tissues of ectodermal origin, data set 2 shows a boundary between cranial mesoderm (CM) and the FMH, and data set 3 shows a boundary separating the Midbrain and the Hindbrain. The formation of this last boundary is a particularly well studied [39], as it plays a crucial role in the development of the brain, the boundary functioning both as a signaling center, also known as the isthmus organizer, and as a physical barrier for the developing brain ventricles [40].

To determine the locus of the physical boundary between tissue compartments, we used the results of our joint image-based force inference (Figure 2A-C) and spatial transcriptomics pipeline (Figure 2D and Figure Sup.2B-C) to obtain an assessment for the statistical likelihood (see Methods). From this approach, it was possible to determine the compartment boundaries for each of the data sets, as shown in Figure 3A-B. Using the data associated with the tension maps shown in Figure 2B, we then computed, for each data set, the homotypic junctional tension for each tissue compartment and the heterotypic junctional tension existing at each boundary. As evidenced by the barplots in Figure 3C, homotypic tensions in tissue compartments are *∼* 12 *−* 35% lower than heterotypic tensions at the compartment boundaries depending on the data set considered, with data set 1 displaying the smallest difference and data set 3 the largest. Taken together, these results seem to rule out a scenario based on DAH or DITH in favour of a mechanism of boundary maintenance based on HIT for all three distinct boundaries.

**Figure 2:**
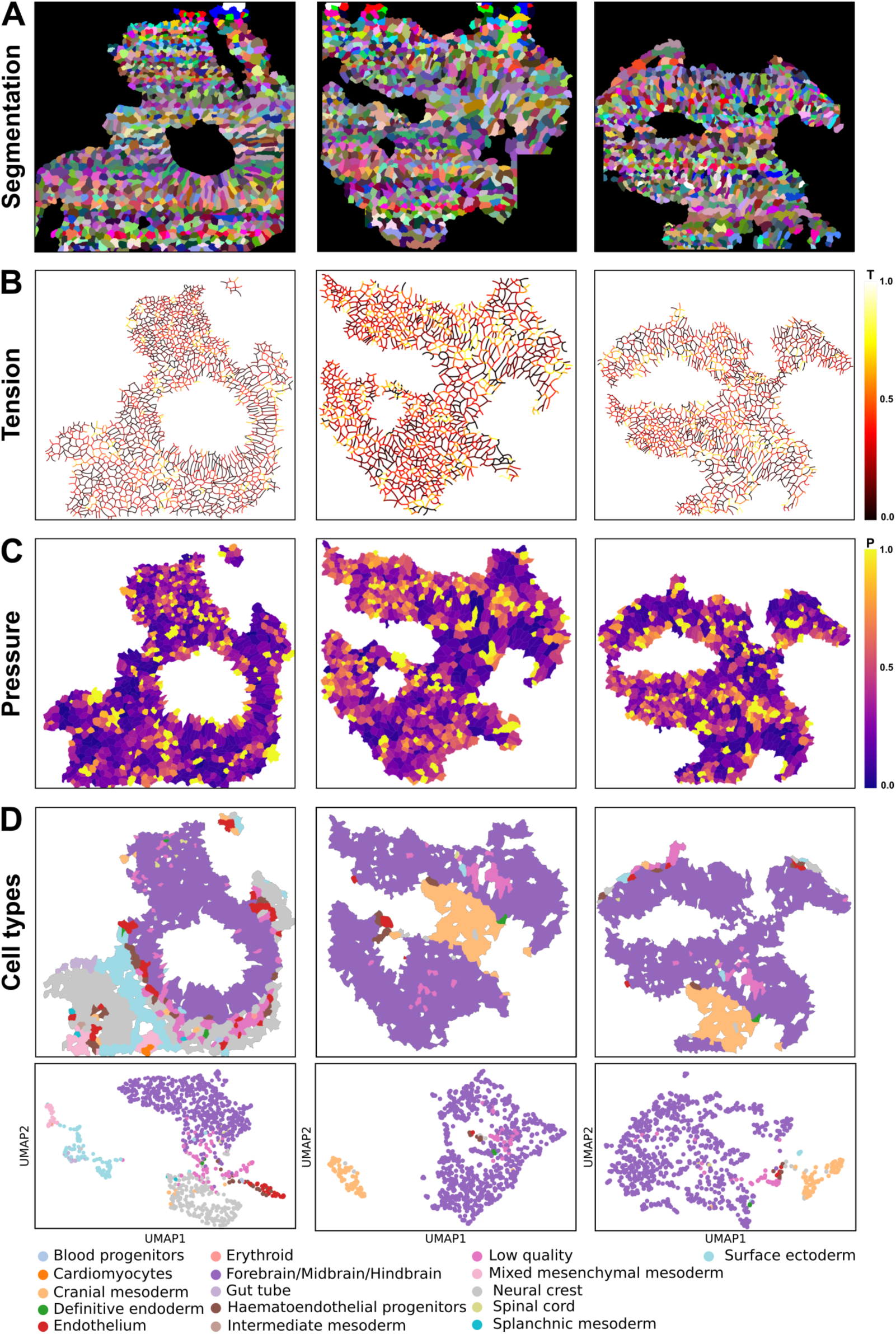
Spatial mechano-transcriptomics profiling of different E8.5 mouse embryo brain regions. **A:** Instance segmentation masks of cell contours (see materials and methods) for data sets 1 (left), 2 (middle) and 3 (right). **B**: Spatial tension maps obtained with the VMSI algorithm from the cell segmentation masks of data sets 1 (left), 2 (middle) and 3 (right). **C** Spatial pressure maps obtained with the VMSI algorithm from the cell segmentation masks of data sets 1, 2 and 3. **D**: Spatial maps and UMAP clustering plots of the cell types present in the data sets 1, 2 and 3. Clusters and cell types were obtained by gene expression analysis on the basis of the seqFISH and inputed gene expression profiles for all cells contained in each data set.

**Figure 3:**
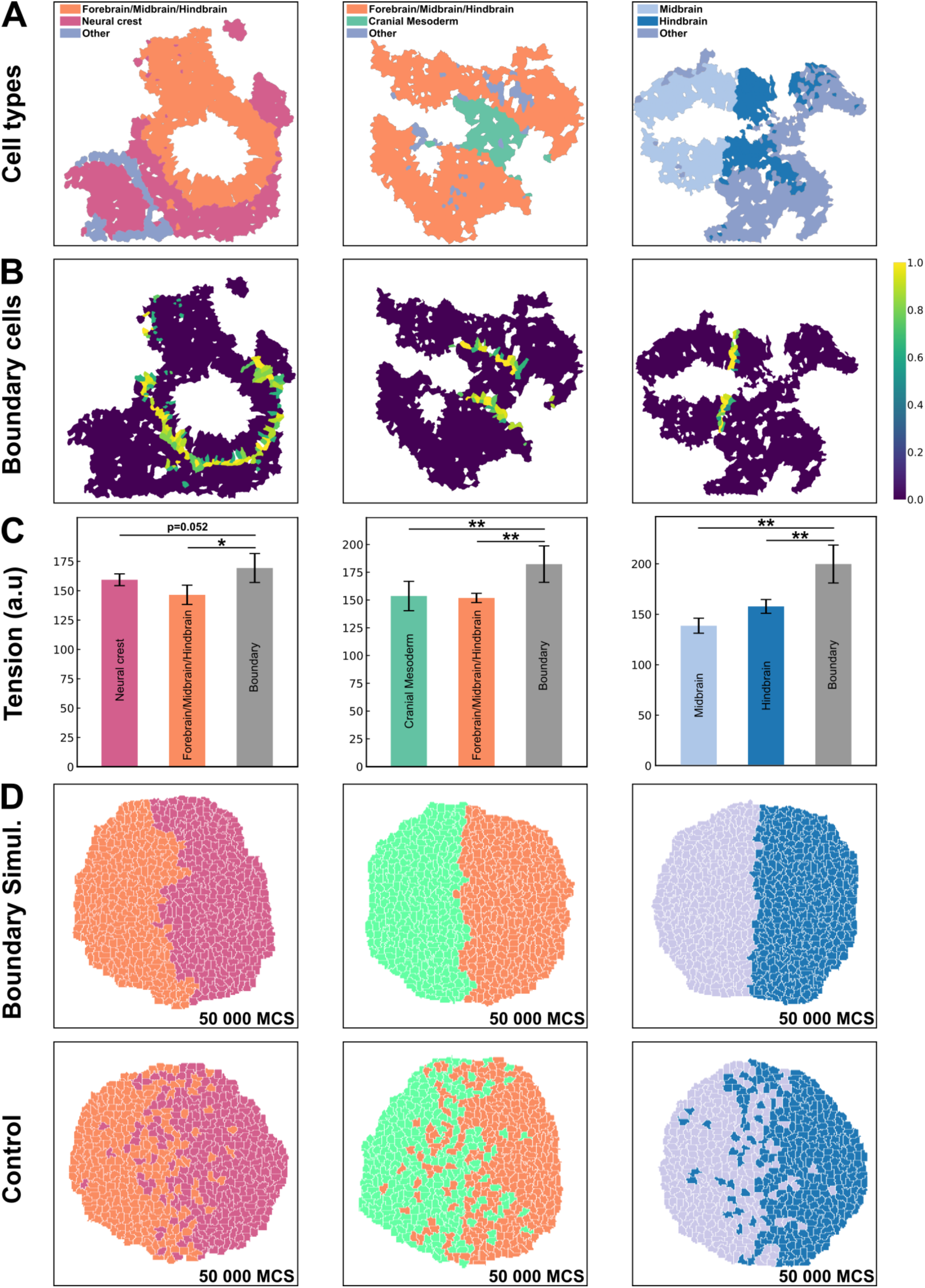
Tissue compartment boundaries defined by gene expression are characterized by a high interfacial tension pattern which is sufficient to explain their maintenance. **A:** Spatial maps of the dominant cell types as defined per gene expression analysis for data sets 1 (left), 2 (middle) and 3 (right). **B**: Spatial maps of the boundary likelihood highlighting cells at the boundary between spatially distinct tissue compartments for data sets 1, 2 and 3. **C**: Bar plots for experimentally measured heterotypic tension (cell-cell tensions for junctions at the boundary between spatially distinct tissue compartments) and homotypic tension (cell-cell tensions for junctions within each tissue compartment) for data sets 1, 2 and 3. Error bars indicate SEM. *^∗^ p <* 0.05 and *^∗∗^ p <* 0.01. Mann-Whitney U test and pair-wise comparisons. **D**: Boundary maintenance simulations based on experimentally measured heterotypic and homotypic tensions for data sets 1, 2 and 3. Top panels display renderings of typical boundary maintenance simulations and bottom panels display renderings of typical control simulations where the homotypic tensions are equal to the heterotypic tensions.

To challenge this scenario, we ran *in silico* experiments using a simple and well characterised biophysical model of multicellular tissues [41, 42]. Specifically, for each data set, we simulated the maintenance of the boundary between the two tissue compartments using experimental values for homotypic and heterotypic tensions with all other model parameters taken as the same. We also ran control *in silico* experiments where the homotypic and heterotypic were taken as equal (see Methods). As shown in the upper panel of Figure 3D, *in silico* experiments confirm that, for all three data sets, a higher interfacial tension at the boundary between tissue compartment is sufficient for boundary maintenance. This phenomenon is characterized by an invariance of the heterotypic boundary length (see Methods) over the length of the simulations (Figure Sup.3D). Here, we also note that the “roughness” of the boundary is inversely proportional to the ratio of the homotypic and heterotypic tensions. Moreover, control simulations confirm that, in the absence of a higher heterotypic tension, cells of both cell types start to mix, leading to a progressive dissolving of the boundary between the tissue compartments, as shown in the lower panel of Figure 3D, and evidenced by the increasing values taken by the heterotypic boundary length in these simulations (Figure Sup.3D). Moreover, further *in silico* simulations using similar numerical parameters, but with different initial conditions, where cells are mixed at random, demonstrated that a higher interfacial tension is also sufficient to explain the formation of segregated tissue compartments via a cell sorting mechanism, as shown in Figure Sup.3B-C.

Overall, HIT appears to be a particularly robust mechanism for tissue compartment boundary maintenance, as even a difference as small as *∼* 10% between homotypic and heterotypic tensions appear to be enough to maintain a boundary. Moreover, spatial tension profiles might provide a highly accurate way to determine with sub-cellular resolution the location of the boundary between tissue compartments. For example, the 1D tension profile at the boundary between the Midbrain and Hindbrain is shown in Figure Sup.4A plotted against the 1D gene expression profiles of Otx2 and Gbx2, two well-characterized markers of the mesencephalon/prosencephalon, and of the rhombencephalon, respectively (Figure Sup.4B). In this case, the position of the boundary can be very accurately pinpointed as the maximum of the 1D tension profile and corresponds to the intersection of the midpoints of the Otx2 and Gbx2 gradients. A similar phenomenon is observed at the boundary between the cranial mesoderm and the FMH tissue compartments, as shown in Figure 4A, where the maximum of the 1D tension profile coincides with the intersection of the midpoints of the Wnt5b and Bmp4 gradients, two well-characterized markers of the FMH and CM respectively (Figure 4B).

**Figure 4:**
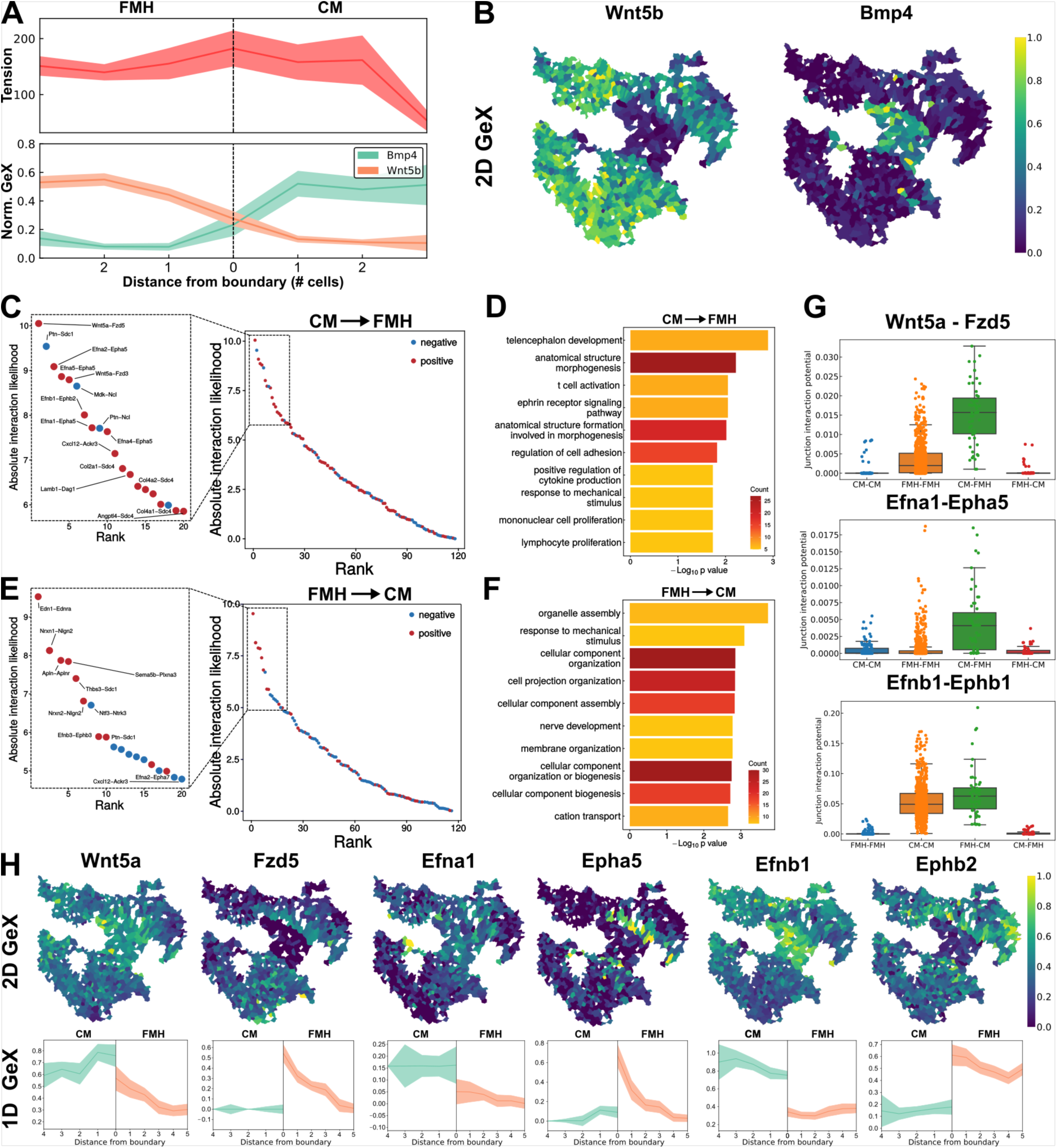
Spatial transcriptomics based ligand-receptor analysis demonstrates ephrin signaling is a molecular determinant of elevated interfacial tension at tissue compartment boundaries. **A:** (Top) 1D tension profile along the tissue compartment boundary for data set 2. (Bottom) 1D gene expression profile for cranial mesoderm (CM) and Forebrain/Midbrain/Hindbrain (FMH) markers Bmp4 and Wnt5b for data set 2 (Mean+/-95%CI). **B**: Spatial gene expression maps for CM and FMH markers Bmp4 and Wnt5b. **C**: Absolute interaction likelihood of LR pairs for pair of cells at the boundary between CM and FMH in data set 2, with ligands expressed in CM cells and receptors in FMH cells. Insert panel diplays top 20 LR pairs with the highest absolute interaction likelihood. **D**: Gene Ontology (GO) enrichment term analysis for the top 100 LR pairs for pair of cells at the boundary between CM and FMH in data set 2, with ligands expressed in CM cells and receptors in FMH cells. GO term are ranked by p-value and gene count. **E**: Same as panel C but for ligands expressed in FMH cells and receptors in CM cells. **F**: Same as panel D but for ligands expressed in FMH cells and receptors in CM cells. **G**: Boxplots and scatter plots of the junction interaction potentials for some top-ranked LR pairs (Wnt5a-Fzd5, Efna1-Epha5, Efnb1-Ephb1). **H**: (Top) Spatial gene expression maps for the same selected LR pairs. (Bottom) 1D gene expression profiles along the tissue compartments boundary for the same LR pairs.

### Ligand receptor analysis identifies putative molecular determinants of elevated interfacial tension at tissue compartment boundaries

Next, we quantified the interaction between transcriptional profiling data and force inference readouts. As higher interfacial tension is a likely physical determinant of tissue compartment boundaries maintenance, we questioned whether the spatial transcriptomics data can provide insight into the molecular mechanisms underpinning this phenotype. As a first step, we used unbiased ligand-receptor (LR) analysis using the spatial gene expression data making use of the CellChatDB LR annotation database (see Methods). We analyzed a dataset involving the boundary between the cranial mesoderm and the FMH tissue compartments, and a dataset involving the boundary between midbrain and hindbrain tissue compartments.

Focusing on cells sharing heterotypic contacts (i.e. on the boundary), it was possible to screen for the expression levels of known LR pairs, and then to compute from their interaction potential (Figure 4 G and Figure Sup.4 G) an absolute interaction likelihood (Figure 4 C, E and Figure Sup.4 C,E), distinguishing the directionality of interactions. Considering only LR pairs displaying a positive interaction likelihood and filtering out the top 50 pairs, we ran a gene ontology (GO) over-representation analysis, whose results are reported in Figure 4 D,F and in Figure Sup.4 D,F. The results emphasize the role of LR signaling in controlling mechanobiological processes such as “response to mechanical stimulus”, “regulation of cell adhesion”, “anatomical structure morphogenesis” and “ephrin receptor signaling pathway” at the tissue compartment boundaries in both data sets.

Notably, considering LR pairs displaying the highest positive interaction likelihoods, it is apparent that some of these pairs involve canonical transmembrane receptors and diffusible ligands such as Wnt5a-Fzd5, which is known to play a crucial role in anterior-posterior axis formation and patterning during mammalian development [43], Fgf18-Fgfr1, which is known to play a key role in the establishment of the boundary between midbrain and hindbrain in mouse [44], and Edn1-Ednra, which has been shown to be a key determinant of cranio-facial morphogenesis in mouse and human [45]. Interestingly, 2D maps and 1D spatial gene expression profiles in Figure 4 H and Figure Sup.4 H show that these LR pairs are involved in directional signaling. For example, in data set 2, the CM acts as an almost spatially homogeneous source of Wnt5a, whereas its expression sharply decreases into the FMH region beyond the compartment boundary. This expression profile is mirrored by the spatial expression pattern of the receptor, Fzd5, which is not expressed in CM, but displays a spatially graded profile in the FMH, with the highest point of the gradient found in cells proximate to the boundary on the FMH side.

Furthermore, a significant fraction of the top LR pairs are ephrin ligand (Efn) receptor (Eph) pairs, such as Efna1-Epha5, Efnb1-Ephb1 or Efnb3-Ephb2, as shown in Figure 4 C,E and Figure Sup.4 C,E. Ephrin ligands are membrane-bound proteins, which can only interact with ephrin receptors expressed in neighbouring cells, with cells expressing a ligand usually down-regulating the expression of its associated ephrin receptor(s) and vice versa [34, 46]. This leads to a characteristic spatial expression pattern, which can be observed in the 2D maps and 1D spatial gene expression profiles in Figure 4 H where the ephrin ligand (Efna1 or Efnb1) is strongly expressed in one of the two tissue compartments (here the CM), while the receptor (Epha5 or Ephb1) is expressed almost exclusively in cells proximate to the boundary in the other tissue compartment (here the FMH). The same characteristic spatial pattern is also observed in data set 3, where one can observe in Figure Sup.4 H the mutually exclusive spatial pattern of Efnb3 and Ephb2 at the midbrain-hindbrain boundary.

Ephrin LR signaling is well known to generate “repulsion” at heterotypic cell-cell contacts and tissue compartment boundaries via downstream signaling pathways that increase interfacial tension for cell-cell junctions located on the boundary [28, 29, 46]. Consequently, the presence of multiple ephrin LR pairs with high interaction likelihood on the boundary between CM and FMH provides a potential mechanistic explanation for the observed higher heterotypic interfacial tension at the boundary, and could be generalised to explain the higher interfacial tension also observed for other tissue compartment boundaries in data set 3 or data set 1.

While these findings emerge naturally from the combined transcriptomic and force inference analysis of the E8.5 mouse embryo, this mechanism constitutes a ubiquitous feature of boundary formation in vertebrates, and has been observed in a variety of developmental contexts such as the boundary between mesoderm and ectoderm in the *Xenopus Laevis* embryo [35], the boundary between the different segments of the hindbrain (rhombomeres) in zebrafish and chick embryos [47], and the boundaries between somites [26, 48], and compartments of the neural tube [49, 50] in zebrafish embryos. In all these systems, the mechanism driving the increase in interfacial tension at the boundary appears to be caused both by an increase in actomyosin contractility, due to myosin II phosphorylation directly downstream ephrin LR signaling via Ephexin mediated RhoA activation, and a localised decrease in cell-cell adhesion, due to selective expression of cell-cell adhesion molecules such as cadherins or protocadherins [51, 46, 29, 50].

While our approach, based on spatial transcriptomics, does not allow us to directly quantify actomyosin activity, we could nonetheless investigate the spatial patterns of cell-cell adhesion molecules at the boundary between CM and FMH in data set 2, as shown in Figure Sup.8 A. Interestingly, CM and FMH display reciprocal patterns of cadherin expression so that when one particular cadherin is up-regulated in one tissue compartment, such as Cdh2 in the FMH or Cdh11 in the CM, it is down-regulated in the other compartment. As homophilic cadherin adhesion is energetically favourable over (or equivalent to, for type I cadherins) heterophilic cadherin adhesion [50], this creates a situation where cell-cell adhesion is markedly decreased at the boundary between tissue compartments and increased within the respective tissue compartment, correlating once again with the pattern of higher heterotypic tension at the boundary and lower homotypic tension within tissue compartments. Previous work suggests that, during zebrafish neural tube compartmentalisation, this mechanism is also regulated via a signaling gradient of the morphogen Shh to Cdh2 and Cdh11 via protocadherin Pcdh19 [49], an observation we are able to corroborate in our system as shown in Figure Sup.8A.

Interestingly, another study on mouse neural tube patterning has shown that a dorso-ventral (DV) gradient of mechanical forces exists in the embryo and leads to a graded activation of YAP signaling along the DV axis, causing a spatially compartmentalized expression of the transcription factor Foxa2 and its downstream transcriptional target Shh [52]. Since such a gradient of mechanical tension exists in the vicinity of the boundary between the CM and FMH in data set 2 (Figure 4 A), it is tempting to speculate that it could also lead to the formation of a gradient of YAP signaling activity in this system, and thus be the origin of the observed Shh gradient at the compartment boundary. This hypothesis is supported by the observation of a graded expression of Cyr61, a well-characterized transcriptional target of Yap, and of Foxa2 and its transcriptional targets, such as Ptch1, at the border between CM and FMH, as shown in Figure Sup.8 B. In addition, as shown in Figure Sup.8 C, markers of neural tube DV patterning, Nkx2.2, Nkx6.1, Pax7, and Pax3, are also expressed at the boundary between the CM and FHM tissue compartments in a spatial sequence that follows the spatial gradient of mechanical forces and is reminiscent of that observed in the mouse neural tube [52].

Overall, these results provide a rational molecular mechanism to explain the higher heterotypic interfacial tension observed at the boundary between tissue compartments in our different data sets and support the conclusion that this mechanism may play an important role in maintaining a sharp boundary at the interface of two tissue compartments. At the same time, this analysis illustrates how the combination of force inference analysis with spatial molecular profiling can provide insight into the mechanism of boundary formation in the context of embryonic development.

### Geoadditive structural equation models detect gene expression modules associated with cellular mechanics while controlling for spatial confounders

Our previous analysis identified putative mechanisms for cooperativity between gene expression and cellular mechanics in establishing and maintaining boundaries during development. To identify additional developmental processes in which cellular mechanical and transcriptional states are coordinated, we next sought to perform unsupervised tests for associations between gene expression and mechanical measurements.

We first tested the association between gene expression and mechanical state for data sets 1 and 2 by using a linear model to regress single-cell gene expression levels on two mechanical quantities, cellular pressure and the magnitude of the cellular stress tensor (see Methods). Statistical analysis identified a number of “mechano-associated” genes, i.e. genes whose expression is significantly up- or down-regulated with cellular pressure or stress tensor magnitude. To identify genes that are confidently associated with mechanics, we searched for genes that were significantly associated with mechanical quantities in both data sets. Figure Sup.7 A shows that there were 150 pressure-associated and 1049 stress-tensor-associated genes shared by data sets 1 and 2, respectively. Among these, a total of 131 mechano-associated genes showed significant association with both pressure and stress tensor magnitude for both datasets. GO overrepresentation analysis (see Methods) showed that this gene set is enriched in genes associated with “cell migration”, “tissue morphogenesis” and “ECM organization” (Figure Sup.7 B), processes that are highly dependent on cellular mechanical state.

To further identify specific signaling pathways and mechanisms, we examined the specific genes involved. Volcano plots in Figure Sup.7 C show these mechano-associated genes for data set 2, highlighting in red some of the 131 top associated genes discussed above. Some genes, such as Hpln1 or Col4a1, are associated with extracellular matrix structure and mechanical properties, while others, such as Ccnl2, are involved in cell cycle regulation or cell metabolism, including Igf2. Some genes such as Arhgef15 (involved in ephrin-LR signaling), Actb, Dchz1 and Rhod are involved in cytoskeleton organization and contractility. Consistently, others are known transcriptional targets of well-characterized mechanotransducers such as Cav1 and Cyr61, which are downstream of Yap. Interestingly, all of the aforementioned genes have gene expression patterns that negatively correlate with the magnitude of the pressure and stress tensor, i.e. they tend to be up-regulated in cells under tensile stress and down-regulated in cells under more compressive stress.

However, a limitation of linear regression testing is that it does not account for spatial confounding effects. Spatial confounding could interfere with the estimated effects because both morpho-mechanical measurements and transcriptomic states are themselves spatially dependent; in particular, tissue regions comprising common cell types or subtypes may show similarities in both bulk mechanical properties and transcriptomic states. Therefore, we performed a second analysis, utilising a geoadditive structural equation model (gSEM), which accounts for spatial confounding effects in both predictor and response variables by modelling and subtracting the spatial confounding effects from both variables, resulting in *spatially regressed* variables with no spatial confounding. This methodology provides a means for rigorously accounting for spatial confounding effects in our data.

We tested the association between gene expression and mechanical state for all 3 data sets using a linear model to regress spatially regressed single-cell gene expression levels on the two spatially regressed mechanical quantities (see Methods). We identified a number of mechano-associated genes; as expected, accounting for spatial confounding resulted in fewer statistically significant genes being identified. Most of these genes appear to be cell type and tissue specific, suggesting that the effects of cellular mechanics on gene expression are context dependent; this highlights the utility of our approach to infer mechanical properties and gene expression in the same cells. Despite differences in specific mechanosensitive genes, GO overrepresentation analysis (see Methods) showed that GO terms relevant to both developmental processes and cellular mechanics were enriched across multiple data sets (Figure 5 B,D and Sup.5 B,D). For example, we identified terms such as “negative regulation of substrate adhesion-dependent cell spreading” and “negative regulation of cell morphogenesis involved in differentiation” enriched in data set 2, while data set 3 was enriched in the terms “regulation of actin cytoskeleton organisation” and “leukocyte migration”.

**Figure 5:**
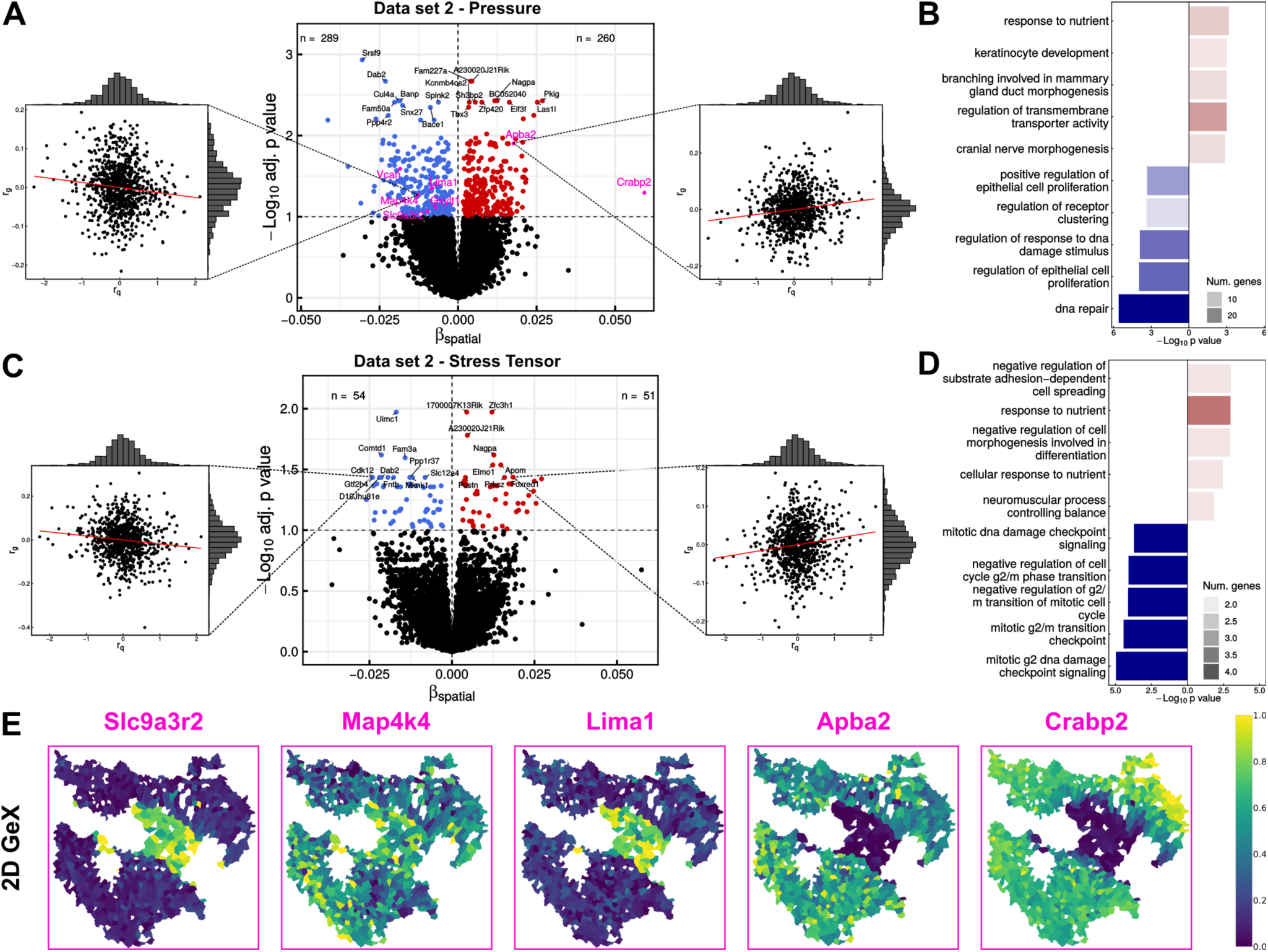
Structural equation regression identifies significant correlations between gene expression and cellular mechanics accounting of spatial confounding effects. **A:** Volcano plot for data set 2 showing for each gene the adjusted p-value (y-axis) plotted against the regression coefficient, *β_spatial_*, obtained by regressing the spatially regressed residual of gene expression on the spatially regressed residual of cellular pressure (x-axis). Side plots represent such linear regressions for two example genes whose spatially regressed gene expression residuals are respectively negatively (left) and positively (right) associated with the spatially regressed residual of cellular pressure. **B**: GO overrepresentation analysis for up- and down-regulated genes with cellular pressure. GO terms are ranked by p-value and gene count. **C**: Volcano plot for data set 2 showing for each gene the adjusted p-value (y-axis) plotted against the regression coefficient, *β_spatial_*, obtained by regressing the spatially regressed residual of gene expression on the spatially regressed residual of the magnitude of the cellular stress tensor. Side plots represent such linear regressions for two example genes whose spatially regressed gene expression residuals are respectively negatively (left) and positively (right) associated with the spatially regressed residual of the cellular stress tensor. **D**: GO overrepresentation analysis for up- and down-regulated genes with cellular stress tensor. GO terms are ranked by p-value and gene count. **E**: Spatial gene expression maps for selected genes (labelled in purple in panel A) displaying significant correlations between gene expression and cellular pressure in both the linear and the structural equation regression analyses.

Volcano plots in Figure 5 A, C and in Figure Sup.5 A, B, show the mechano-associated genes identified for data sets 2 and 3. We found that, although there was generally a low degree of overlap between genes identified as significantly associated in the linear regression analysis above and the gSEM analysis, the inferred effect sizes showed good correlation across both analyses (Figure Sup.5 C, E). Furthermore, several genes were highlighted in both analyses. Many of these genes have known roles in regulating cellular mechanical properties, for example Slc9a3r2 (NHERF2), Lima1 and Crabp2 (Figure 5 E). Slc9a3r2 interacts with and regulates the ERM complex, which couples the actomyosin cortex with the cell membrane and enables forces generated through cytoskeletal dynamics to influence the overall mechanical properties of the cell and more particularly the cell-cell junctional tensions [53]. Lima1 is also relevant in actin cytoskeletal dynamics through regulating actin fibre crosslinking and depolymerisation [54], while Crabp2, a component of the retinoic acid signalling pathway, has previously been shown to modulate mechanosensing in the context of pancreatic cancer [55]. Our analysis also revealed a number of novel links between mechanics and gene expression. One such example is Apba2, which interacts with and stabilizes the amyloid precursor protein (APP). Interestingly, previous work has shown that aggregation of the amyloid-*β* peptide generated by APP affects the mechanical properties of single cells in a pathological context [56]. This novel association suggests a potential role for Apba2, and thus APP, in responding to changes in mechanical state during development.

### Analysis of nonlinear associations between gene expression and mechanical properties identifies distinct patterns of association with cellular mechanics

We next turned to investigate non-linear associations between cellular mechanics and gene expression at the single cell level. To that aim, we ranked cells in each data set by either cellular pressure or stress tensor magnitude, and computed smoothed expression value estimates using a local weighted median metric. Subsequently, we used scHOT [57] to identify statistically significant patterns of association between the weighted median gene expression and cellular mechanical property. Significant gene-mechanics associations were then clustered using hierarchical clustering to identify clusters of genes with consistent association patterns. We performed this analysis for both data set 2 (Figure 6) and data set 3 (Figure Sup.6). For data set 2, we obtained 7 clusters of genes associated with pressure and 4 clusters of genes associated with stress tensor magnitude. For data set 3, we obtained 7 clusters of genes associated with pressure and 5 clusters of genes associated with stress tensor magnitude.

**Figure 6:**
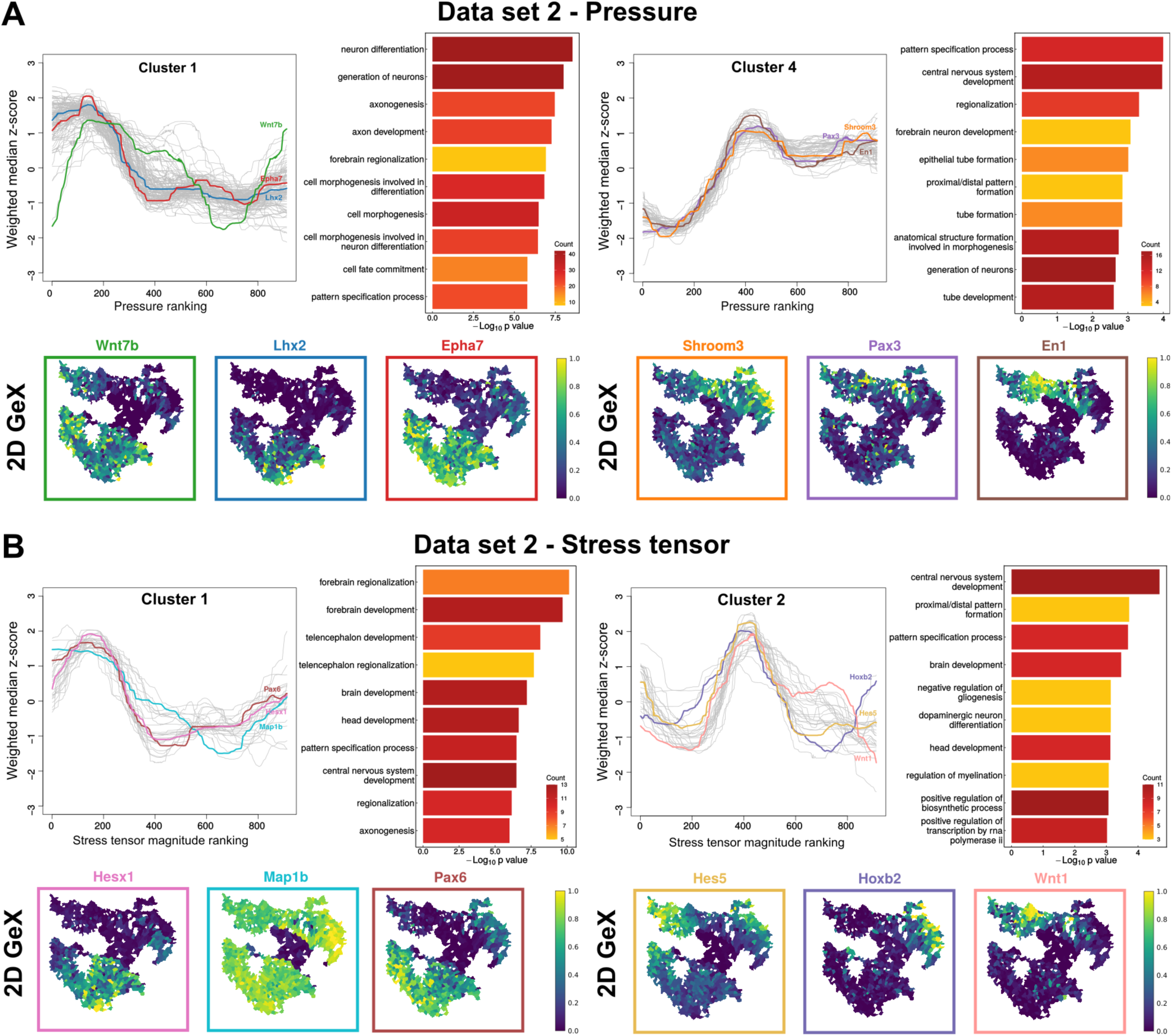
Statistical analysis of higher-order interactions establishes the existence of significant non-linear associations between gene expression and cellular mechanics. **A:** Analysis of nonlinear associations between gene expression and cellular pressure in data set 2. The summary statistic used is the weighted median gene expression. For each gene, the association between this statistic and the cellular pressure ranking is tested. Significant (p.adj ¡ 0.1) association profiles are Z-normalised and clustered. Line plots of the weighted-median expression Z-score against cellular pressure ranking are shown for selected clusters, along with bar plots showing GO overrepresentation analysis of genes in each cluster. Lower panels display spatial gene expression maps for example genes with representative behaviours. GO terms are ranked by p-value and gene count. **B**: Analysis of nonlinear associations between gene expression and cellular stress tensor magnitude in data set 2. Analysis was performed as for (A) above. Line plots of the weighted-median expression Z-score against cellular pressure ranking are shown for selected clusters, along with bar plots showing GO overrepresentation analysis of genes in each cluster. Lower panels display spatial gene expression maps for example genes with representative behaviours. GO terms are ranked by p-value and gene count.

The clusters identified in data set 2 revealed that different clusters showed distinct patterns of association with cellular mechanics, and different spatially localised patterns of expression, suggesting that mechanical differences between tissue regions may influence regions-specific gene expression. Interestingly, we also identified functional differences between genes in different clusters. In data set 2, amongst genes non-linearly associated with pressure, cluster 1 displayed a sigmoid expression profile where gene expression is up-regulated at low intracellular pressure and down-regulated after a certain pressure threshold (Figure 6A). GO overrepresentation analysis (see Methods) revealed that these genes were involved in developmental processes such as “cell fate commitment”, “neuron differentiation” and “glial cell migration”. Genes in cluster 4 display the opposite behavior, being expressed at a low levels prior to becoming up-regulated at higher intracellular pressure when values exceeded a certain threshold (Figure 6A). These genes were found to be associated with a variety of cellular and developmental processes such as “pattern specification process”, “epithelial tube formation”, and “forebrain neuron development”. Reflecting the GO overrepresentation analysis, we also observed known master regulators of neural development (e.g. Wnt7b, Lhx2, Pax3, En1), as well as genes involved in cell adhesion and contractility (e.g. Epha7, Shroom3) within the same clusters, suggesting cooperativity between cellular mechanics and regulation of developmental processes.

As for genes non-linearly associated with stress tensor magnitude, clusters 1 and 4 also displayed the two kinds of sigmoid response previously encountered (Figure 6B). Genes in cluster 1 were associated with “forebrain development” and “telencephalon development”, and were up-regulated at low stress tensor magnitude prior to sharply decreasing their expression beyond a certain threshold. Mirroring this behaviour were genes of cluster 2, which were associated with “central nervous system development” and “proximal/distal pattern formation”, and were down-regulated at low and high stress tensor magnitude, with expression within only a narrow range of stress tensor magnitude values. Notably, the expression profiles displayed by gene clusters 1 and 2 showed a remarkable sensitivity, suggesting that the expression of these genes is regulated by either a mechanosensitive band-pass (cluster 2) or band-stop (cluster 1) filter. Corroborating this, we observed similar band-pass behaviour in gene clusters identified in data set 3 (Figure Sup.6); again, we also observed co-localisation of factors important in development and regulators of cellular mechanics within the same clusters. This suggests that these band-pass and band-stop behaviours may be general mechanisms for coupling mechanics and gene expression during development. While such non-linear gene expression dependencies have been engineered in synthetic bacterial and mammalian systems in response to external biochemical signals [58], their observation in the setting of a native tissue is, to our knowledge, unprecedented.

## Discussion

In this study, we presented a first computational framework for combined spatial transcriptomics and image-based mechanical force inference at single cell resolution. Using synthetically generated images of multicellular tissues, we showed that our approach is accurate and robust to noise associated with confocal fluorescence imaging of immunostained tissue sections and cell instance segmentation. We demonstrated that our framework can be applied to *in situ* hybridization-based spatial transcriptomics datasets by performing an integrated analysis of a seqFISH data set of the E8.5 mouse embryo. Using three different brain regions from two different embryos as benchmark datasets, we were able to perform an integrated analysis of mechanical forces and gene expression at single cell resolution.

Our analysis revealed that tissue compartment boundaries predicted by differential gene expression analysis are associated with specific spatial patterns of increased cell-cell junctional tension. Using a biophysical modeling approach, we showed that elevated heterotypic interfacial tension is sufficient to explain the long-term maintenance of boundaries between adjacent differentiated tissue compartments. Furthermore, biophysical simulations of systems initialized with randomly mixed cell types suggested that high heterotypic tension, as opposed to homotypic tension, may also be responsible for the initial establishment of these boundaries. To infer the underlying molecular determinants of this mechanical phenotype, we used ligand-receptor analysis to show that ephrin ligand-receptor signaling most likely regulates the elevated tension at tissue compartment boundaries through localized increased actomyosin contractility and differential cell-cell adhesion.

Finally, using our framework, we were able to statistically quantify the relationship between mechanical forces and gene expression patterns at single cell resolution. A geoadditive structural equation model revealed a significant number of genes whose expression was either positively or negatively associated with mechanical forces. Interestingly, these gene sets were associated with a variety of cellular and tissue processes such as cell migration, cell metabolism, mechano-transduction, response to morphogens and hormones, or tissue morphogenesis. Furthermore, higher-order analysis of the association between single-cell gene expression and cellular mechanics revealed the existence of several families of mechano-associated behaviors. Some genes were up-regulated at lower cellular pressures and down-regulated at higher ones, and vice versa. Notably, the expression of some genes was found to be up- or down-regulated over a narrow range of mechanical forces, suggesting the existence of mechanosensitive band-pass and band-stop filters.

However, our analysis highlights the limitations of the seqFISH technology. First, the fidelity of this approach is highly dependent on the quality of staining and 2D sectioning. The quality of membrane immunostaining can hinder the segmentation of individual cell contours, leading to inaccurate recovery of cell junction curvatures, imprecise inference of mechanical forces, and difficulties in processing large data sets. Alternative membrane staining strategies, such as the use of antibodies against other membrane proteins or against other components of the cell membrane such as glycolipids [59], can improve membrane staining and allow large-scale automated cell segmentation and accurate mechanical force inference.

Second, our current approach focuses on 2D slices. While it has been shown that 2D force inference is a good proxy for 3D inference for simple isotropic cellular ensembles, such as those found in components of early mouse or nematode embryos [60], this is not generally true for non-planar and anisotropic systems. For example, the seqFISH data set used in this analysis includes whole-embryo sagittal sections, where some regions may intersect the plane of the section rather than being parallel to it. This means that the inferred 2D stress tensor captures only a subset of the information present in the full 3D stress state of a cell. In addition, E8.5 mouse embryos contain a variety of regions that are not populated by cells, but by extracellular matrix (ECM) and fluid-filled cavities whose mechanical properties influence the mechanical behavior of adjacent tissue layers in ways that cannot be captured by the present two-dimensional method. Generalizing the current framework through three-dimensional gene expression profiling [61], cell segmentation, and force inference [62] will be a critical step toward a more integrative and precise understanding of the reciprocal role of mechanical forces and gene expression, cell fate decisions, and tissue morphogenesis during development. Taking advantage of improved staining and 3D imaging, future studies will aim to extend the scope of our analysis by incorporating additional morphometric measures to capture cell shape, such as point cloud-based methods [16] or Fourier shape descriptors [15]. In addition, the measurement of additional genomic modalities, such as metabolomics, proteomics, and chromatin accessibility, as well as metrics that capture the nature of the local cell environment, such as the size and composition of the cell neighborhood or the coarse-grained stress tensor, could help us to better understand how cellular mechanical and transcriptional phenotypes are regulated and integrated at the tissue and organismal level.

Overall, our computational framework can be applied directly to ISH-based spatial transcriptomics datasets with minimal additional processing required. Although some previous studies have performed combined analysis of single-cell morphometrics and gene expression [63], and others have investigated the relationship between mechanical forces or mechanical properties and expression of individual genes [64, 65], integration of mechanical force inference and spatial transcriptomics at single-cell resolution has not been previously reported. The work presented here contributes to our understanding of the interplay between mechanical forces and gene expression at the cell and tissue level and provides an innovative and powerful tool that can be applied to other spatial transcriptomics datasets to further investigate this interplay in a variety of physiological and pathological contexts.

## Methods

### Transcriptomics quantification

A previously published multi-embryo seqFISH data set was used to examine the utility of the spatial mechano-transcriptomics workflow [24]. In this approach, the abundance and positions of individual transcripts were obtained at subcellular resolution for 387 genes across sections of three mouse embryos at developmental stage E8.5. This data set was used to impute a broader pattern of gene expression taking advantage of the mouse gastrulation atlas data set, a previous single-cell atlas obtained from scRNA-seq analysis using a 10X Genomics pipeline [66]. We targeted the correlation between cell mechanics and gene expression in the context of boundary formation in three different brain regions (Figure 1B), spanning the intersection between the Forebrain/Midbrain/Hindbrain (FMH) and Neural Crest (NC), data set 1, the boundary between the Cranial Mesoderm (CM) and FHM, data set 2, and an upper brain region involving the Midbrain/Hindbrain boundary (MHB), data set 3.

#### Image segmentation

High quality segmentation masks are essential for accurate image-based mechanical force inference. As the existing segmentation masks for the E8.5 mouse seqFISH data set exhibited high variability across biological regions and replicates with frequent instances of over- or under-segmentation, we reprocessed the imaging data sets as follows: We first performed automated segmentation of DAPI labeled cell nuclei using a custom deep learning pipeline. The ground truth data set used for training was composed of 12 image and mask pairs tilled into 16 random 256×256 pixel image patches and split into 3 batches comprising training, validation, and test data sets in a 70:15:15 ratio. The convolutional neural network which was trained for binary segmentation involved a custom “light weight” U-Net with a reduced depth of one level as compared with the original implementation [67] resulting in a network with *∼* 0.5 million nodes and using ELU instead of ReLu as activation functions. Training was carried out using Tensorflow 2.0 and Keras 2.8 libraries [68], using a custom loss function combining weighted binary cross-entropy and dice index loss, and using the Adam optimizer, a batch size of 16 and a learning rate of 0.0001. Then, the resulting nuclei centroids were used as seeds to a initialise a watershed algorithm [69] to generate cell instance segmentation masks on the basis of the averaged E-Cadherin and *β*-Catenin immunostaining fluorescence signals. The cell contour segmentation masks were further pre-processed and curved edges between cell-cell contacts were identified via circular arc fitting. Poor-quality edges were manually corrected using Fiji [70].

#### Circular arc polygon tiling

Following image segmentation, circular arcs approximating the locus of cell boundaries and their contact points are required for downstream stress inferences [71]. This results in a circular arc polygon (CAP) tiling. More precisely, the CAP tiling fits a circular arc parameterised by the centre of curvature ***ρ****_αβ_* and radius of curvature *R_αβ_* to each cell-cell junction between two cells *α, β*. In cases where the cell-cell junction is not curved, or exhibits inconsistent curvature (e.g., ‘wiggly’ boundaries where the sign of curvature changes along the boundary), a straight line was fit to the junction instead. The curve-fitting procedure, as well as the criteria for identifying straight junctions, were adapted from [71].

#### Spatial transcriptomics processing

Cells identified in the corrected segmentation were correlated with cells in the original segmentation using a pairwise Jaccard index. Real overlaps were defined as cells with greater than 0.1 Jaccard similarity, and all overlaps were filtered out. Weights for each cell in the original segmentation mask for each cell in the corrected segmentation mask were calculated using the fraction of overlap in the segmentation masks. Cells in the corrected segmentation with *≤* 0.4 total overlap were filtered out. The resulting weights were used to compute corrected expression matrices, using a weighted mean of both the imputed expression values and raw counts for genes profiled by seqFISH. Corrected raw counts were further normalised by the total mRNAs identified in each cell, and log-transformed.

#### Tissue boundaries defined by transcriptomic profiles

Boundaries within the three data sets were defined using a boundary likelihood metric. For a given cell *i* with neighbours *N* and two sets of cell types *A* and *B*, the boundary likelihood between *A* and *B* at cell *i* was defined as:

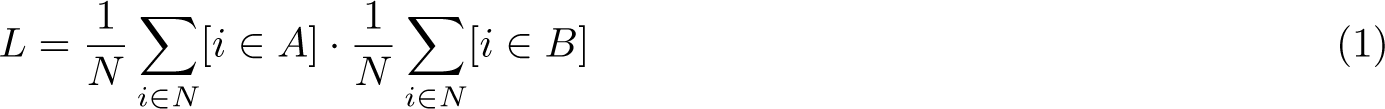

A threshold of *L >* 0.15 was applied to identify cells at a boundary. The boundary within the ‘embryo 2 midbrain-hindbrain’ region was defined manually, similarly to the method applied in the original study [24].

To investigate properties of cell-cell junctions at boundaries, each cell was assigned a distance to boundary *d*, defined as the number of neighbours between that cell and the closest cell belonging to the boundary. A cell-cell junction between the cell pair *{α, β}* was defined as ‘near-boundary’ if *min*(*d_α_, d_β_*) *≤* 5 and ‘at-boundary’ if *min*(*d_α_, d_β_*) = 0. At-boundary junctions were then classified as homotypic if both cells belonged to the same cell type set, or heterotypic otherwise.

#### Ligand-receptor signalling analysis across tissue compartment boundaries

Log-transformed, normalised imputed gene expression values were derived after correction using the method described above and used for analysis of ligand-receptor signalling potential across tissue boundaries.

Ligand-receptor annotations from the CellChat database [72] were obtained with Omnipath [73] and filtered for ligand-receptor pairs for which both ligand and receptor showed non-zero expression in our transcriptomic data. An ‘interaction potential’ *P_L→R,α,β_* = *L_α_ · R_β_* was defined for each ligand-receptor pair *{L, R}* across the cell pair *{α, β}* to quantify the potential degree of signalling through the receptor. This definition takes into account the directionality of signalling interactions and allows for the signalling through the receptor to be investigated independently for both tissues at a boundary.

Ligand-receptor signalling interactions were compared for two spatially adjacent cell types *{A, B}* using the interaction likelihood metric *l*, defined as:

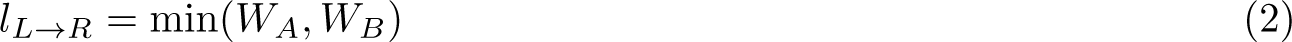

where *W_A_* represents the Wilcoxon rank-sum test statistic between the interaction potential distribution *P_L→R,α,β_* for *α ∈ A, β ∈ B* and the interaction potential distribution for *{α, β} ∈ A*. Signalling interactions were ranked by interaction likelihood, with negative interaction likelihoods (i.e., where the *{A, A}* interaction likelihood or *{B, B}* interaction potential is higher than the *{A, B}* interaction likelihood) filtered out, and genes in the top 50 interactions were tested for over-representation of GO terms compared to the total set of ligand and receptor genes (see the GO analysis section).

### Mechanics quantification

#### Inferring tension from images

There are a variety of methods for inferring inter-cellular stress in tissues at mechanical equilibrium, i.e., where the tensions at each vertex of the cell array sum to zero [23]. These methods vary in sophistication, which mechanical features are inferred, and dependence on the image segmentation quality. At the most basic level, segmentation-free methods exploit the correlation between cell shape anisotropy and stress anisotropy to derive coarse-grained estimates of tissue stress in tissues where accurate cell segmentation cannot be performed [74]. If segmentation is possible, but there is high noise that prevents a precise determination of the geometry of cell-cell junctions and vertices, methods such as chord inference can be used that model cell-cell junctions as straight lines and therefore discount the contribution of cell pressure to the geometry of the cell array [20]. Tangent inference methods improve on chord inference by using the angle between cell-cell junctions at vertices. This allows for less noisy output but requires more precise image segmentation. However, cell pressures are again not taken into account in this approach [75]. Recent methods are able to infer both cell junction tension and cell pressure by measuring the curvature at cell-cell junctions as well as vertex angles. However, these methods generally require increased segmentation precision and are not robust to noise. The VMSI method [22] circumvents these issues by inferring both pressures and tensions simultaneously from fitted CAP tilings instead of the segmented image, as the CAP tiling provides additional noise reduction over the segmentation itself. Hence, we build on and extend the VMSI method to probe the mechanical properties of seqFISH generated data.

#### Mechanical phenotypes

Following [71], three mechanical phenotypes were computed for each cell *α* and each adjacent cell pair (*α*, *β*): the cellular pressure *p_α_*, the cell-cell junctional tension *T_α,β_*, and the stress tensor *σ_α_*.

Given a CAP tiling, let *ρ_αβ_* be the center of the curvature of the circular arc at the (*α, β*) cell-cell junction, and let *R_αβ_* be the radius of curvature of the same arc. Force balance equations result in geometrical constraint variables *{q_α_, θ_α_}*, which parameterize the curvature center and radius:

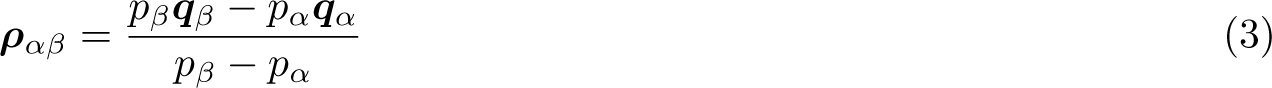

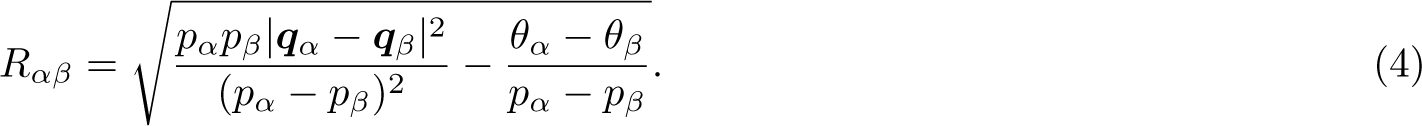

##### Cellular pressure

Cellular pressures were computed in two steps [71]. First, initial values for cell pressures *p_α_* and geometric constraint parameters *q* enforced the condition that *ρ_αβ_*, the centre of curvature to the edge vertices *r_i_, r_j_*, must be perpendicular to the edge tangents *τ_i_, τ_j_*, minimizing the functional:

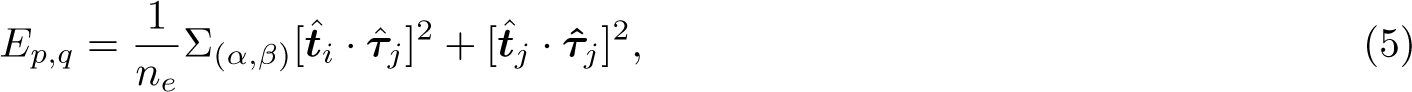

where *t*^^^*_i_* is the unit vector along *r_i_ − ρ_αβ_* and *τ*^*_i_* is the edge tangent at vertex *i*. Similarly, initial values *θ* optimized the functional:

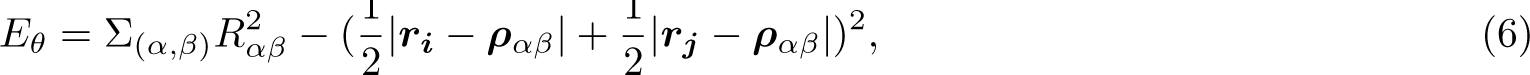

where *ρ_αβ_* were calculated using the *{p, q}* values determined previously. Second, the initial values (*p_α_, q_α_, θ_α_*) were used to instantiate the gradient descent optimization of the objective

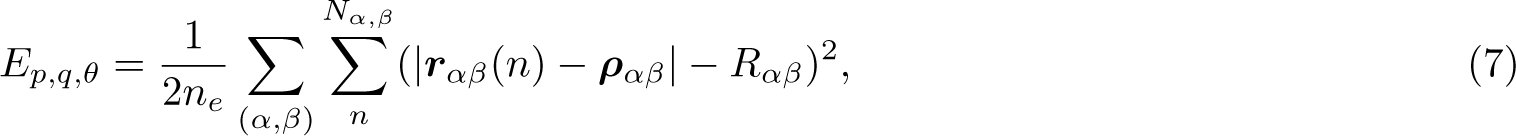

finding the mechanical equilibrium parameters resulting in a CAP tiling which best approximated the one obtained through image segmentation. Here, *r_αβ_*(*n*) denotes the *n^th^* pixel along the circular arc approximation of the edge between cells (*α, β*) in the segmented CAP tiling, and *n_e_* denotes the total number of edges.

##### Cell-cell junctional tension

Cell-cell junctional tensions were computed as functions of the corresponding cellular pressures and the corresponding radius of curvature using the Young-Laplace law: *T_αβ_* = (*p_α_ − p_β_*)*R_αβ_*.

##### Stress tensor

2D cellular stress tensors *σ_α_* were defined from the inferred cellular pressures and cell-cell junctional tensions using Batchelor’s formula: [76]

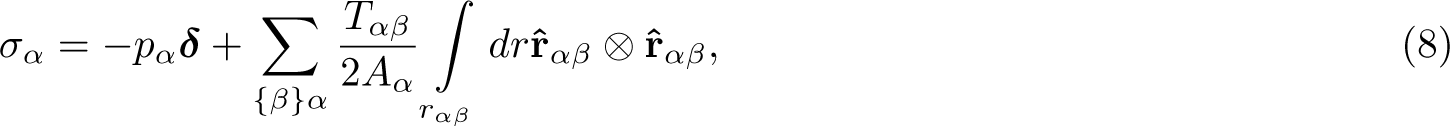

where *p_α_* and *A_α_* denote the pressure and area of cell *α* respectively, *T_αβ_* denotes the junctional tension between adjacent cells (*α, β*), and *r*^*_αβ_* is the unit vector along the junction. The resulting 2×2 stress tensor encodes all of the stress information of a cell [60]. Using the elements of the cellular stress tensor, five interpretable descriptors of the mechanical state of a cell can be computed: the two eigenvalues of the stress tensor, the stress tensor magnitude, the stress tensor anisotropy and the stress tensor orientation. The stress tensor magnitude was defined as the sum of its eigenvalues, its anisotropy was defined as the eccentricity of ellipse formed by the two eigenvectors and its orientation was defined as the angle between the major axis of the ellipse formed by the two eigenvectors and the x-axis of the image.

### Practical considerations

#### Calibration via mask processing

For force inference results to be valid, variational methods such as VMSI [71] assume that all vertices between cells are threefold, as a large proportion of vertices with more than three cells would violate the assumption of mechanical equilibrium [23]. The dual triangulation used by VMSI explicitly forbids fourfold (or greater) vertices. In our implementation (Figure 1 A), these vertices are filtered prior to inference by recursively splitting each invalid vertex into two vertices in the direction of greatest variance of neighbour vertices until all vertices are threefold. Further, VMSI assumes that all angles between cell junctions at a vertex are convex. Concave vertices under the VMSI formulation imply negative tension at one of the junctions [23], a situation that is hard to motivate biologically and beyond the scope of the method. Therefore, concave vertices are assumed to be a precision error in the cell segmentation, and are dealt with by moving the vertex until all angles between junctions are concave, as shown in Figure 1A.

#### Robustness checks via simulations

Synthetic images of 2D multicellular tissues (Figure Sup.1A) for which the ground-truth values of cell pressures and cell-cell junction tensions are known were generated to test the accuracy and robustness of force inference. The estimated force inference values were highly correlated with their corresponding ground truth values (Spearman’s *ρ >* 0.96, Figure Sup.1B) across a range of average pressure differentials (Sup.1 F and G) and image sizes (Sup.1E). Furthermore, our approach showed robustness against noise in the measured vertex position as well as occasional incorrect merging of adjacent cells (under-segmentation) during image segmentation (Sup.1 C and D). This is significant as these are common sources of error in instance cell segmentation and demonstrates the practical applicability of our image-based force inference algorithm to real microscopy images.

#### Optimization details

We developed a Python implementation of the VMSI algorithm. All optimization steps were performed using the augmented Lagrangian method with a subsidiary L-BFGS algorithm using the NLopt optimisation library [77]. Analytic Jacobians were supplied for the objective and constraint functions for increased speed and accuracy.

### Tissue boundary maintenance and cell sorting simulations

To simulate tissue compartments boundary maintenance and cell sorting, we used a custom C++ implementation [42] of the Cellular Potts Model (CPM) [41]. In this framework, multicellular tissues are represented as 2D lattices of pixels, *k*, partitioned into *N* cells. Each cell, *i*, is composed of all the lattice sites with a pixel value equal to *i*, with *i ∈ {*1 *. . . N}*. Each cell is assigned a cell-type, *τ_k_*, which is defined at the pixel-level, and is thus a function of its position on the lattice. The dynamics of this system is driven by two components: a membrane surface energy term and an elastic deformation term. The membrane surface energy is determined by cell-cell junctional tensions – the outputs of the mechanical force inference algorithm – and controlled by a cell-type dependent parameter *J_τk,τk′_* . The elastic deformation term enforces the condition that cell volumes *V_i_* do not significantly deviate from a target value *V*_0_ and are parameterized by a bulk modulus *κ*. The cell-cell interactions and the volume constraints can be combined into a global energy function, *E*:

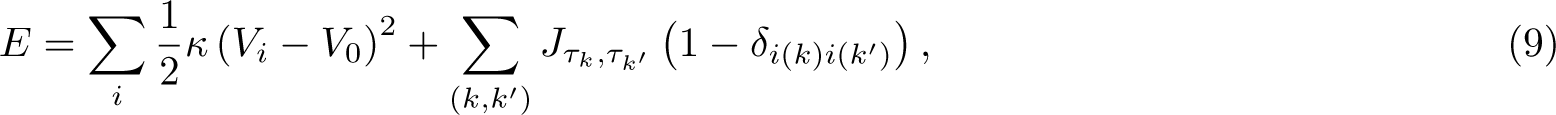

where *i* represents the cell index and (*k, k*) represent pairs of neighbouring pixels. *δ_i_*_(*k*)*i*(*k*_*′*_)_ takes the value of unity when both pixels belong to the same cell and 0 otherwise, in order to solely account for interactions at cell-cell junctions. Moreover, *J_τ,τ ′_* = *J_hom_*, if *k* and *k^′^* belong to two cells of the same cell-type and *J_τ,τ ′_* = *J_het_*, if *k* and *k^′^* belong to two cells of a different cell-type. The system dynamics results from the iterative minimisation of this energy function through the Metropolis Monte-Carlo algorithm [78], where the level of noise in the system is accounted for by a temperature parameter *T* . Time is here expressed in Monte Carlo steps (MCS), where 1 MCS corresponds to an average of one iteration per pixel over the whole lattice. For the simulations described in Figure 3 and Figure Sup.3, parameters are set to *V*_0_ = 40, *κ* = 1.0, and *T* = 10.0. The numerical values used in simulations for parameters *J_hom_* and *J_het_* differ for each data set and are those inferred for the homotypic and heterotypic junctional tensions reported in Figure 3C. For simulating tissue compartment boundary maintenance, a cell aggregate was initially split in two by a straight boundary separating two distinct cell types. For cell sorting simulations, initial conditions were set to a cell aggregate where the two cell types were allocated at random. For all simulations the total number of cells was set to *N* = 540 cells, which were equally assigned to both cell types considered. All simulations were run for 50,000 MCS and at least in 6 replicates. To quantify the boundary maintenance and sorting dynamics, we computed the Heterotypic Boundary Length, *l_HB_*, defined as the total length of the interface between cells of a different cell-type:

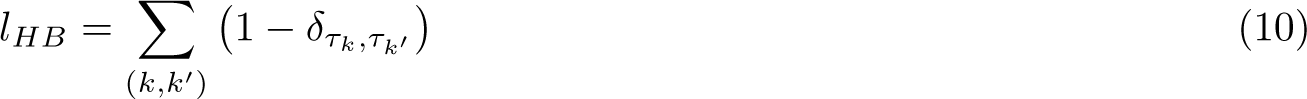

where *δ_τ,τ ′_* takes the value of unity when *k* and *k^′^* belong to cells of the same cell-type and 0 otherwise. As shown in Figure Sup.3 C, *l_HB_* decreases over time in cell sorting simulations, as cells of a different cell-type sort out in spatially distinct clusters. However, as shown in Figure Sup.3 D, during boundary maintenance simulations, *l_HB_* remains constant, as long as the boundary between the two tissue compartments is maintained.

### Statistical mechano-transcriptomics analysis

#### Linear regression

Associations between gene expression and mechanical properties were first tested using linear regression. Two mechanical properties, cell pressure and stress tensor magnitude, were tested. Mechanical properties were first log-normalised, and regression was performed using the linear model:

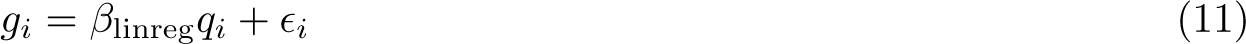

for a gene *g* and log-transformed mechanical property *q* with a standard normal error term *ɛ*. Significance was determined by the FDR-adjusted p-value of the t-test statistic for the regression coefficient *β*_linreg_, using a threshold of *p.adj ≤* 0.05 after correction using the Benjamini-Hochberg procedure.

#### Structural equation regression

The linear regression model described above does not take into account potential spatial confounding effects, which are likely present in our data. Spatial location can influence both gene expression and cell mechanics. On a local scale, cells in close spatial proximity influence each others’ expression profiles and mechanical properties through cell-cell interactions. On a global scale, cell types, which are highly spatially structured, play important roles in dictating gene expression and mechanical properties. To account for this potential spatial confounding in both the predictor and response variables, we therefore used a geoadditive structural equation model (gSEM) [25].

The gSEM accounts for spatial confounding by fitting a thin plate regression spline to determine the effect of spatial location on both the predictor and response variables,

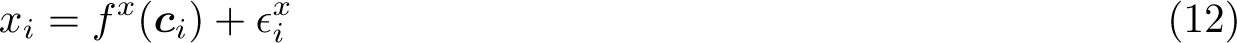

where *x* is the predictor or response variable, *c_i_* is the spatial coordinate associated with *x_i_*, and *ɛ* is a standard normal error term. The fitted values are then subtracted from the predictor and response variables to give the spatially regressed residual *r*. The gSEM is the linear model

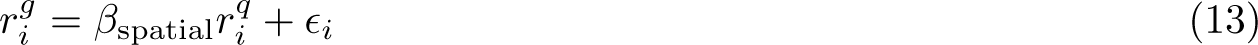

where *r^g^* = *g − f ^g^*(*c*) is the spatially regressed residual for the normalised gene expression *g* and *r^q^* = *q − f ^q^*(*c*) the residual for the log-transformed mechanical property *q*. Significance testing was performed as for the linear regression model described above.

#### Non-linear associations between gene expression and mechanical forces

The regression methods described above uncover linear relationships between gene expression and mechanical forces. However, nonlinear relationships may also exist. Specifically, gene expression may be associated with mechanical stress in a nonlinear monotonic manner, which could indicate the presence of feedback loops or auto-regulation in mechanosensitive signalling pathways. Alternatively, nonlinear non-monotonic associations may suggest the presence of band-pass filter-like mechanisms wherein gene programs are only active within certain ranges of cellular mechanical stress or pressure.

To test this, we ranked cells by pressure or stress tensor magnitude, and computed a smoothed estimate of gene expression along the ranked cells using the weighted median. Given a weighting scheme *w* that assigns a weight to each cell, and a vector of gene expression values *g*, the weighted median is the solution of the optimisation problem

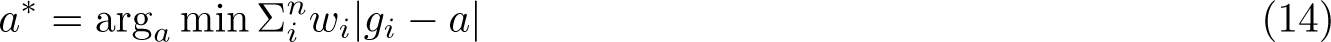

We used a triangular weight matrix with span 0.1, which assigns non-zero weights to cells which have a pressure/stress tensor ranking within 10 percentiles of a given cell. This corresponds to *∼* 150 cells in our data sets.

Significance testing was performed using scHOT [57], which implements a permutation test-based method. scHOT randomly permutes the cell ranking and recomputes the weighted median along the permuted ranking. The variance of the weighted median values was then used as a test statistic. We used 200 permutations per gene to ensure robust significance estimates. Permutation test p-values were then corrected for multiple hypothesis testing using the Benjamini-Hochberg procedure.

To ensure computational tractability, the top 3,000 highly variable genes were identified using Scanpy, and scHOT testing was used to identify genes for which the weighted median expression changes significantly along the pressure or stress tensor magnitude ranking. A threshold of *p.adj ≤* 0.1 was used to determine significantly associated genes. Weighted median profiles were then clustered using hierarchical clustering and the number of clusters was estimated automatically using dynamicTreeCut. Over-representation of GO terms within clusters compared to the total set of scHOT-tested genes was then performed as described in the GO analysis section.

#### Gene Ontology analysis

Gene ontology over-representation analysis was performed using the enrichGO() function from the clusterProfiler R package [79]. Each gene set was tested for over-representation of GO terms against a background set composed of all 29,452 genes for which expression values have been measured or imputed. As GO terms are organised hierarchically, the simplify function was used to remove redundant terms.

## Author contributions

A.H, R.H, B.D. performed image segmentation, computational analysis framework development for mechanical force inference and spatial transcriptomics data, tissue boundary maintenance and sorting simulations. A.H, B.D.S and B.D conceived, managed and supervised the project. All authors wrote and reviewed the manuscript.

## Data availability

Data presented in this study are available from the corresponding authors upon request and will be made openly accessible on public depositories at the time of publication.

## Code availability

A Python implementation of the combined force inference, morphometrics and transcriptomics framework, as well as the code required to reproduce the subsequent analyses presented here, can be found at: https://github.com/ Computational-Morphogenomics-Group/TensionMap/.

## Acknowledgments

The authors would like to thank Nicholas Noll, Nicolas Harmand and Sylvie Hénon for helpful discussions on image-based mechanical force inference and Louis de Benoist for early contributions to the mechanical force inference framework. A.H. gratefully acknowledges the support of the University of Cambridge Herchel Smith Fund through a Herchel Smith Postdoctoral Research Fellowship and the support of Darwin College Cambridge through a Research Fellowship. A.H and B.D.S acknowledges the support of the core funding to the Wellcome / CRUK Gurdon Institute (203144/Z/16/Z and C6946/A24843).

## Supplementary Figures

**Figure Sup.1:**
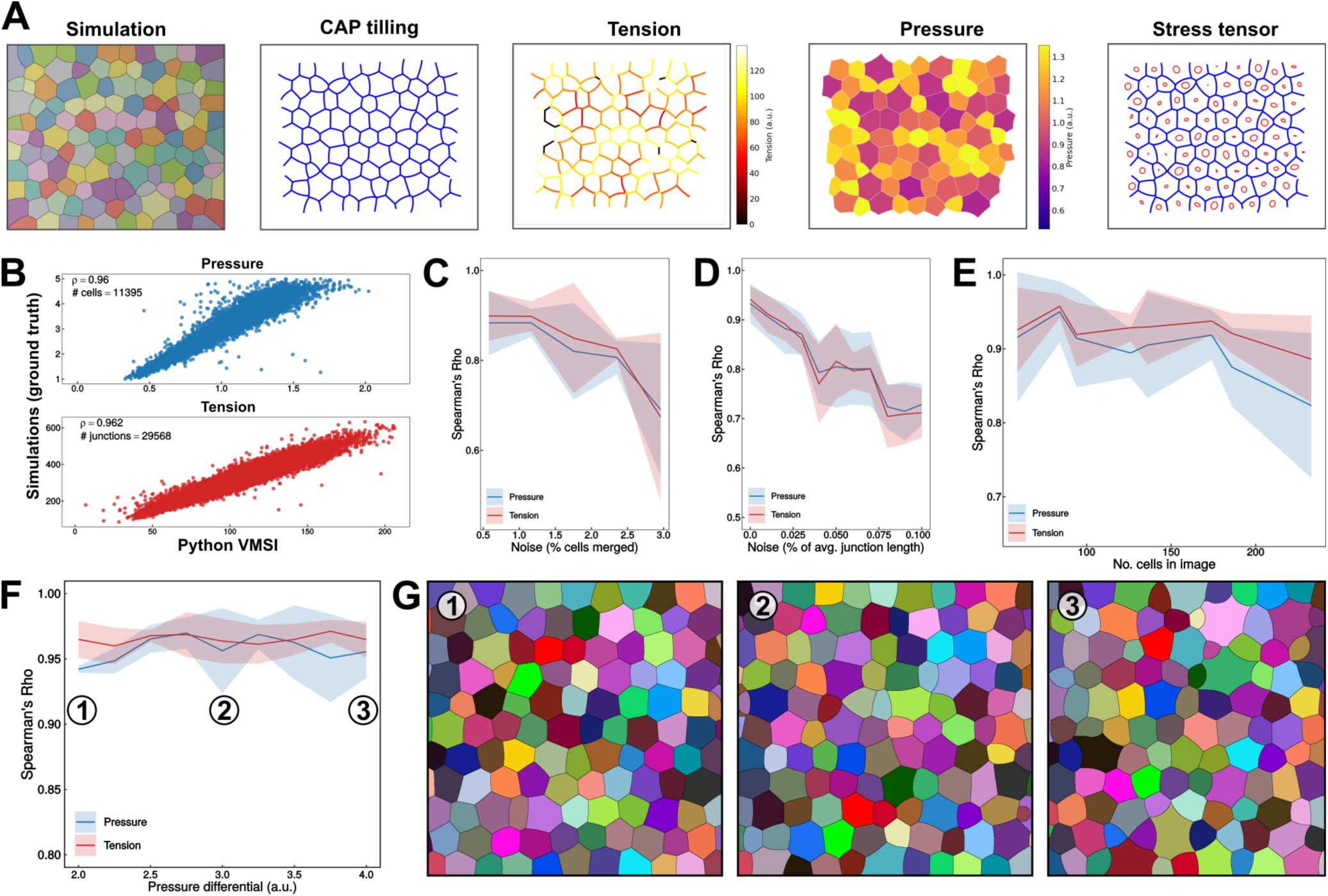
Benchmarking the python VMSI algorithm against ground truth simulated data. **A:** Bench-marking pipeline for our python implementation of the VMSI algorithm. Ground truth simulations for confluent 2D multicellular tissues are generated from known cell-sell tension and cellular pressure distributions (see materials and methods). CAP tillings are obtained from the ground truth simulations and our python implementation of the VMSI algorithm is used to generate cell-cell junction tension maps and cellular pressure maps. Tension and pressure maps are used to compute the cellular stress tensor maps. **B**: Correlation plots representing ground-truth values for cell-cell tensions (bottom) and cellular pressures (top) versus values of the same quantities inferred by our python implementation of the VMSI algorithm. **C**: Correlation between ground truth values and inferred values for cell-cell tensions and cellular pressures as the function of random perturbations to cell segmentation (percentage of merged segmented cells). **D**: Correlation between ground truth values and inferred values for cell-cell tensions and cellular pressures as the function of random perturbations to vertex position (percentage of average junction length). **E**: Correlation between ground truth values and inferred values for cell-cell tensions and cellular pressures as the function of the number of cells in the image. **F**: Correlation between ground truth values and inferred values for cell-cell tensions and cellular pressures as the function of varying cellular pressure differentials. Strength of the correlations were measured using Spearman’s rho coefficient for panels B, C, D, E and F. **G**: Snapshots of ground truth simulations for varying cellular pressure differentials. Each panel is associated with particular data points in panel F.

**Figure Sup.2:**
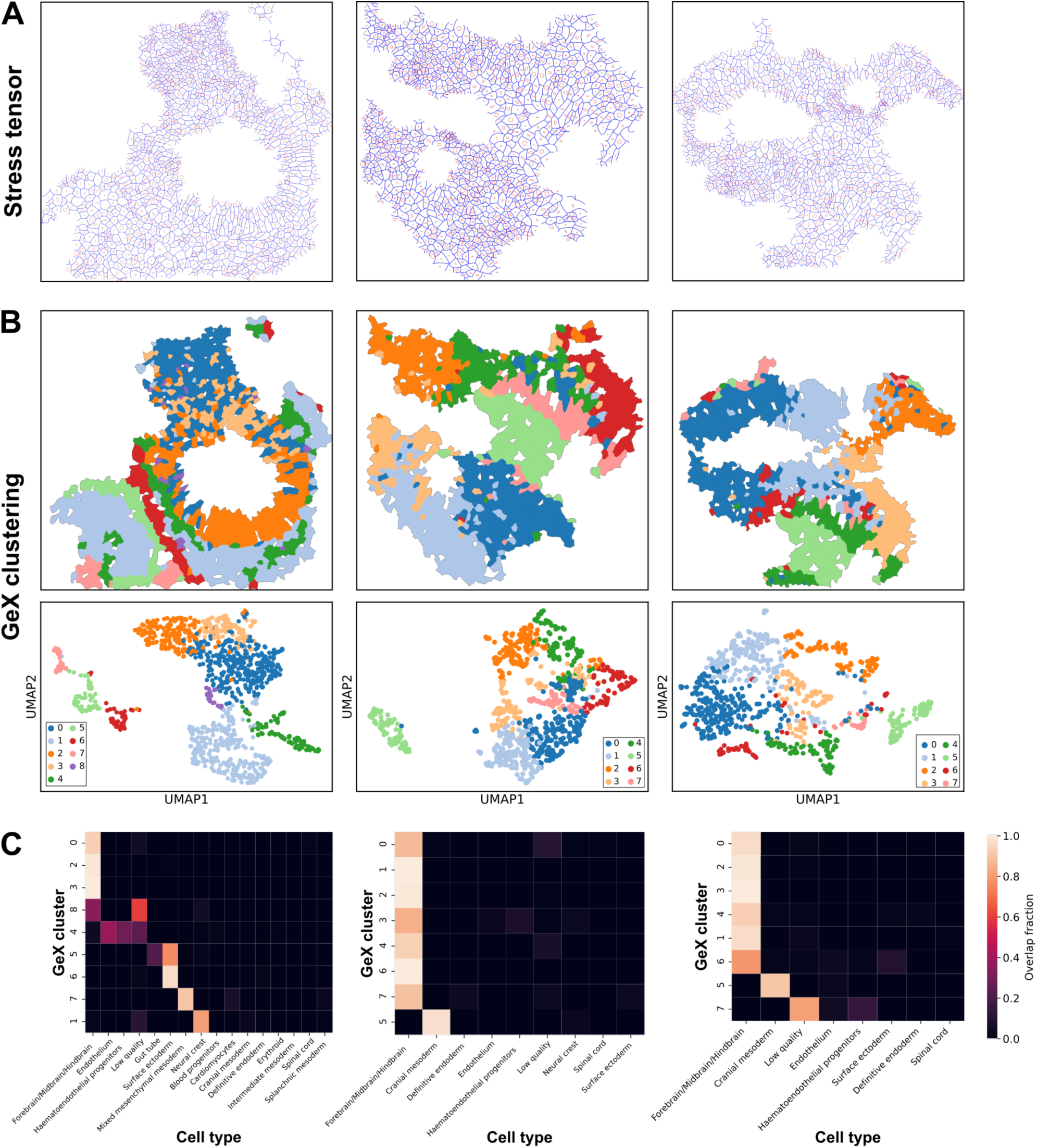
Spatial mechano-transcriptomics profiling of 3 different E8.5 mouse embryo brain regions. **A:** Spatial stress tensor maps computed from the cell-cell tension and cellular pressure maps inferred with the VMSI algorithm on the basis of the cell segmentation masks of data sets 1 (left), 2 (middle) and 3 (right). Each cellular stress tensor is represented by an ellipse whose long and short axis lengths are associated with the eigenvalues of the stress tensor and their orientation with those of the associated eigenvectors. **B**: Spatial maps and UMAP clustering plots of the clusters obtained for data sets 1 (left), 2 (middle) and 3 (right) on the basis of the seqFISH and inputed gene expression profiles for all cells contained in each data set. **C**: Heat maps representing the overlap between the clusters obtained on the basis of gene expression profiles for data sets 1 (left), 2 (middle) and 3 (right), as outlined in panel B, and the cell types, outlined in Figure 2 panel D.

**Figure Sup.3:**
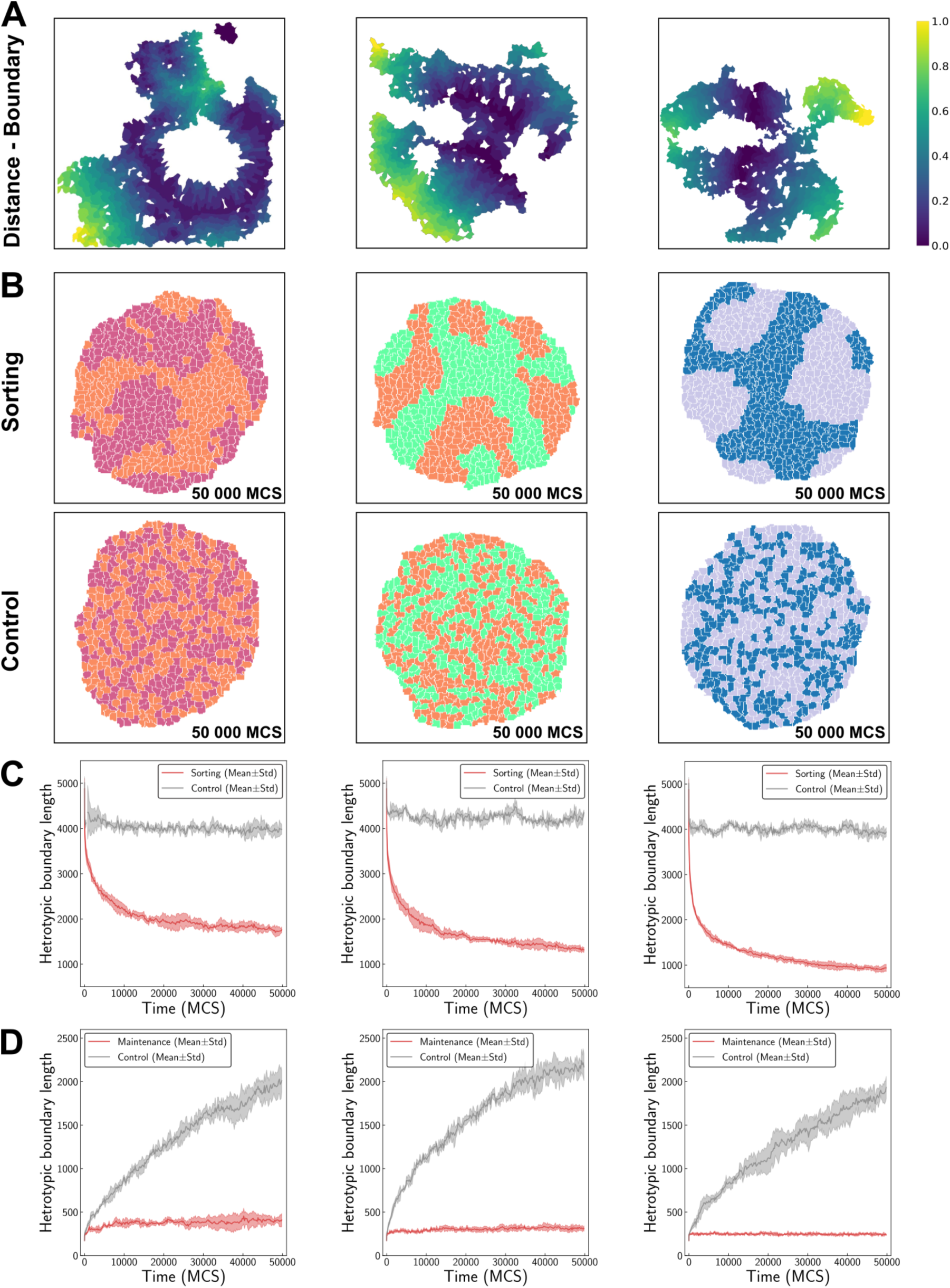
Tissue compartment boundaries defined by gene expression are characterized by a high interfacial tension pattern which is sufficient to explain their maintenance. **A:** Spatial maps of the normalised distance to the tissue compartment boundary for data sets 1 (left), 2 (middle) and 3 (right). **B**: Cell sorting simulations with cells initially mixed at random and using experimentally measured heterotypic and homotypic tensions for data sets 1 (left), 2 (middle) and 3 (right). Top panels display renderings of typical cell sorting simulation and bottom panels display renderings of typical control simulations where homotypic tensions are equal to heterotypic tensions. **C**: Heterotypic boundary length as function of time for cell sorting simulations and associated control simulations for data sets 1 (left), 2 (middle) and 3 (right). **D**: Heterotypic boundary length as function of time for boundary maintenance simulations and associated control simulations for data sets 1 (left), 2 (middle) and 3 (right).

**Figure Sup.4:**
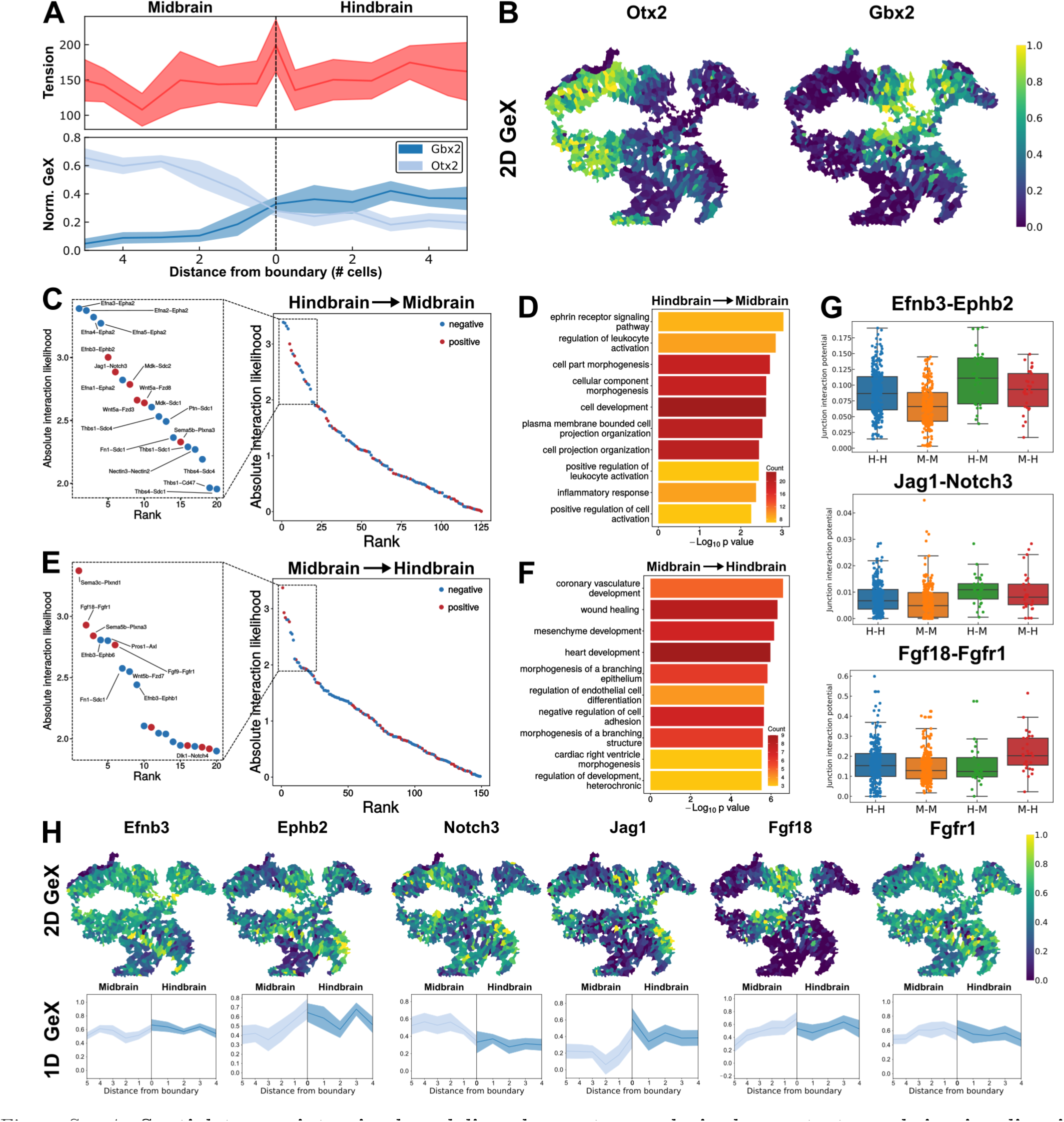
Spatial transcriptomics based ligand-receptor analysis demonstrates ephrin signaling is a molecular determinant of elevated interfacial tension at tissue compartment boundaries. **A:** (Top) 1D tension profile along the tissue compartment boundary for data set 3. (Bottom) 1D gene expression profile for hindbrain and midbrain markers Gbx2 and Otx2 for data set 3 (Mean+/-95%CI). **B**: Spatial gene expression maps for hindbrain and midbrain markers Gbx2 and Otx2. **C**: Absolute interaction likelihood of LR pairs for pair of cells at the boundary between hindbrain and midbrain in data set 3, with ligands expressed in hindbrain cells and receptors in midbrain cells. Insert panel diplays top 20 LR pairs with the highest absolute interaction likelihood. **D**: Gene Ontology (GO) enrichment term analysis for the top 100 LR pairs for pair of cells at the boundary between hindbrain and midbrain in data set 3, with ligands expressed in hindbrain cells and receptors in midbrain cells. GO term are ranked by p-value and gene count. **E**: Same as panel C but for ligands expressed in midbrain cells and receptors in hindbrain cells. **F**: Same as panel D but for ligands expressed in midbrain cells and receptors in hindbrain cells. **G**: Boxplots and scatter plots of the junction interaction potentials for some top-ranked LR pairs (Efnb3-Ephb2, Jag1-Notch3, Fgf18-Fgfr1). **H**: (Top) Spatial gene expression maps for the same selected LR pairs. (Bottom) 1D gene expression profiles along the tissue compartments boundary for the same LR pairs.

**Figure Sup.5:**
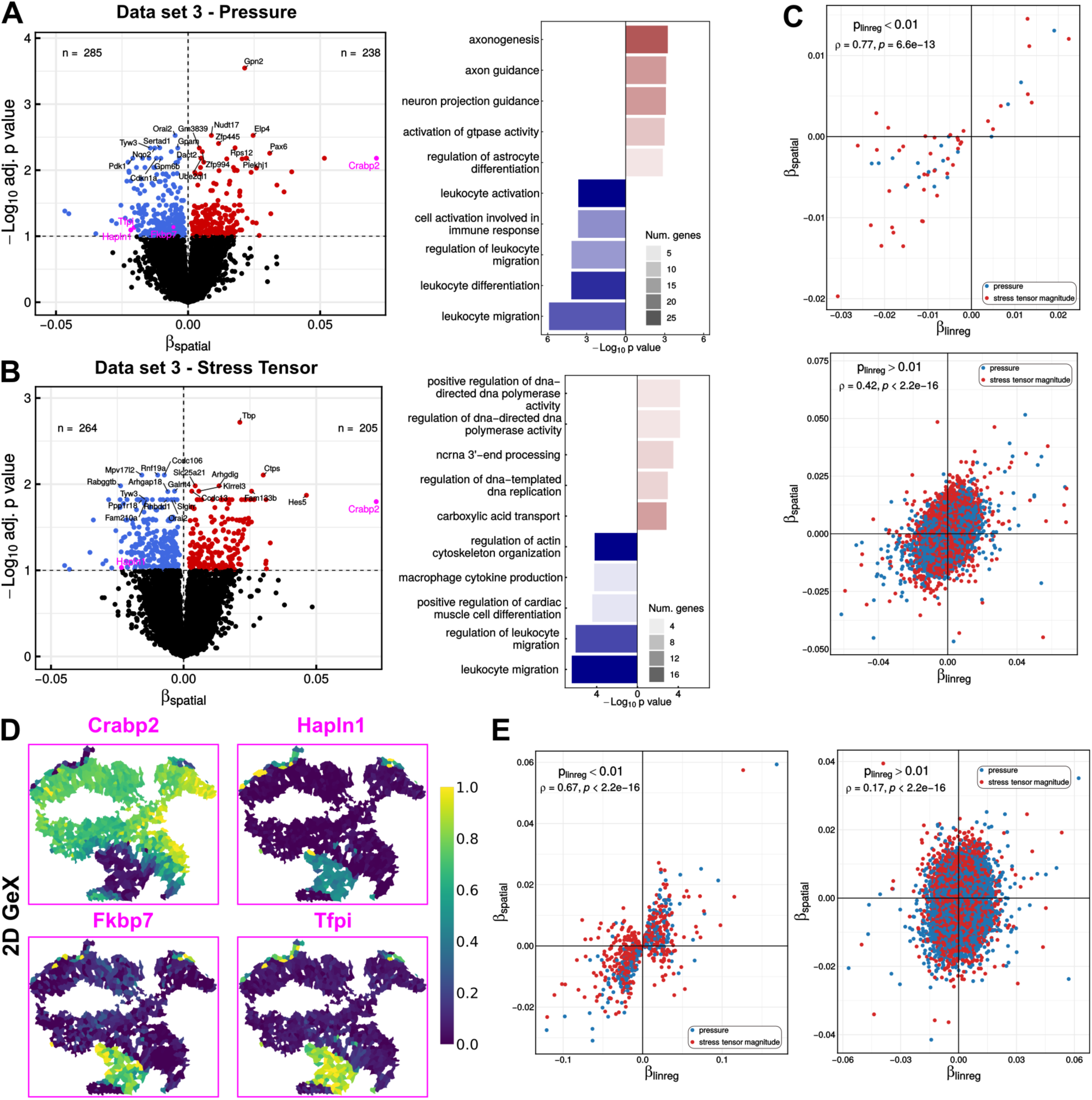
Structural equation regression establishes the existence of significant correlations between gene expression and cellular mechanics independent of any spatial confounding effect. **A:** (Left) Volcano plot for data set 3 representing for each gene the adjusted p-value in respect with the regression coefficient, *β_spatial_*, obtained by structural equation regression of the spatially regressed residual of gene expression as a function of the spatially regressed residual of cellular pressure. (Right) GO enrichment terms analysis for up- and down-regulated genes with cellular pressure. GO terms are ranked by p-value and gene count. **B**: (Left) Volcano plot for data set 3 representing for each gene the adjusted p-value in respect with the regression coefficient, *β_spatial_*, obtained by structural equation regression of the spatially regressed residual of gene expression as a function of the spatially regressed residual of the magnitude of the cellular stress tensor. (Right) GO enrichment terms analysis for up- and down-regulated genes with cellular stress tensor. GO terms are ranked by p-value and gene count. **C**: Correlation plots for *β_spatial_* and *β_linereg_* in data set 3, with *p_linreg_ <* 0.01 (top) and *p_linreg_ >* 0.01 (bottom). **D**: Spatial gene expression maps for some of the genes (labelled in purple in panel A and B) displaying significant correlations between gene expression and cellular pressure in both the linear and the structural equation regression analyses in data set 3. **E**: Correlation plots for *β_spatial_* and *β_linereg_* in data set 2, with *p_linreg_ <* 0.01 (top) and *p_linreg_ >* 0.01 (bottom).

**Figure Sup.6:**
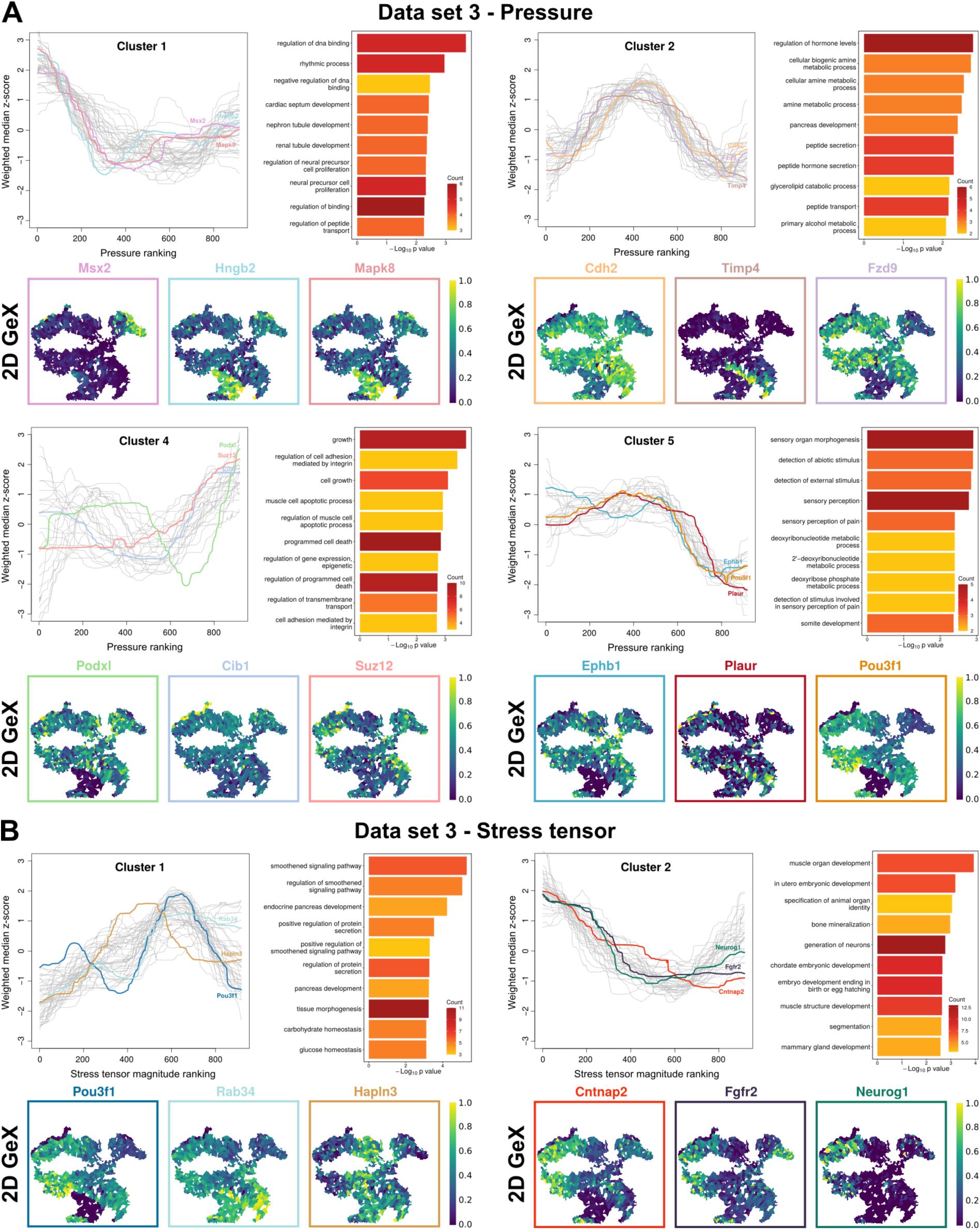
Higher order statistical analysis establishes the existence of significant non-linear correlations between gene expression and cellular mechanics. **A**: Analysis of higher order associations between gene expression and cellular pressure in data set 3 using scHOT. Weighted median of gene expression is plotted as a function of the cellular pressure ranking for genes in clusters 1, 2, 4 and 5 along with associated GO overrepresentation analysis. Lower panels display spatial gene expression maps for example genes representative of each cluster. GO terms are ranked by p-value and gene count. **B**: Analysis of higher order associations between gene expression and cellular stress tensor magnitude in data set 3 using scHOT. Weighted median gene expression is plotted as a function of the cellular stress tensor magnitude for genes in clusters 1 and 2, along with associated GO overrepresentation analysis. Lower panels display spatial gene expression maps for some example genes representative of each cluster. GO terms are ranked by p-value and gene count.

**Figure Sup.7:**
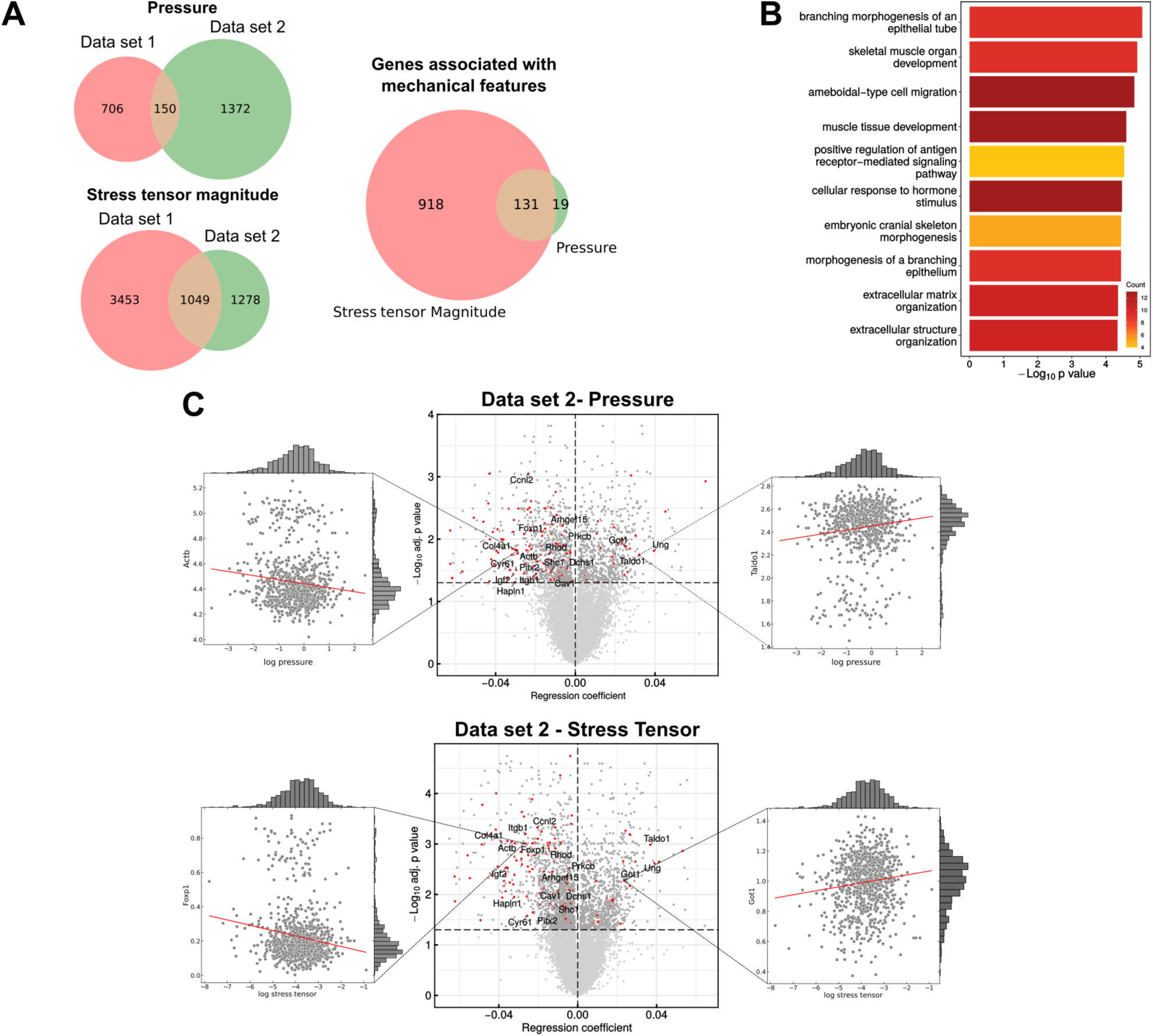
Linear regression analysis establishes the existence of significant correlations between gene expression patterns and cellular mechanical states. **A:** Venn diagrams showing for data set 1, data set 2, and their overlap, the number of genes which expression patterns display statistically significant correlations with cellular pressure (top), stress tensor magnitude (bottom) and both mechanical quantities (right). **B**: Gene Ontology (GO) enrichment term analysis for the 105 genes whose expression pattern show significant correlation with both cellular pressure and stress tensor magnitude. GO term are ranked by p-value and gene count. **C**: (Top) Pressure volcano plot for data set 2 representing for each gene the adjusted p-value in respect with the regression coefficient obtained by linear regression of the gene expression levels as a function of the cellular pressure. Side plots represent such linear regressions for two example genes respectively negatively (left) and positively (right) correlated with cellular pressure. (Bottom) Stress tensor magnitude volcano plot for data set 2 representing for each gene the adjusted p-value in respect with the regression coefficient obtained by linear regression of the gene expression level as a function of the magnitude of the cellular stress tensor. Side plots represent such linear regressions for two example genes respectively negatively (left) and positively (right) correlated with the magnitude of the cellular stress tensor.

**Figure Sup.8:**
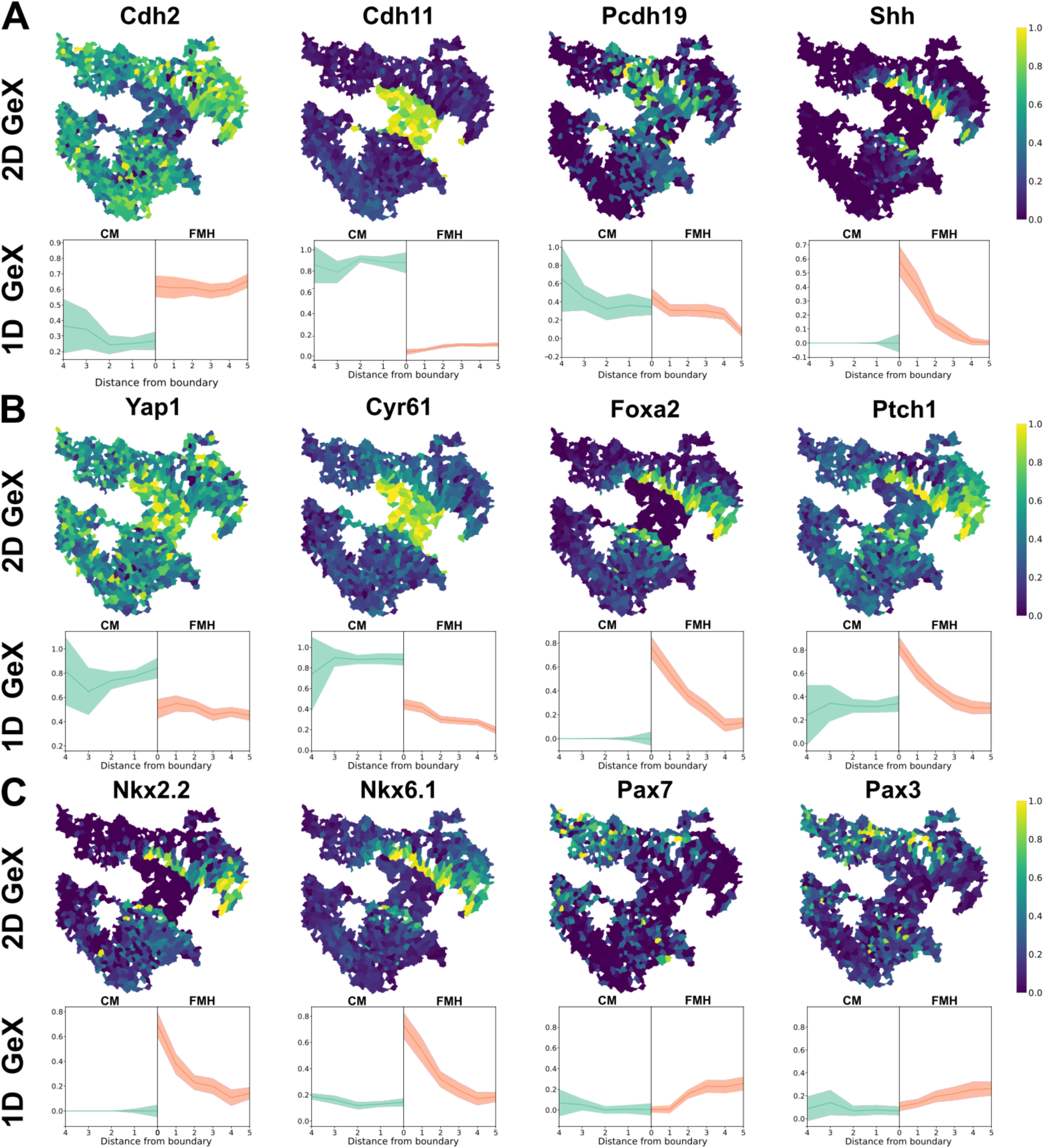
Spatial expression patterns of mechano-associated genes and some of their transcriptional targets at the boundary between CM and FMH in data set 2. **A:** (Top) Spatial gene expression maps for Cdh2, Cdh11, Pdch19 and Shh. (Bottom) 1D gene expression profiles along the tissue compartments boundary for the same transcripts. **B:** (Top) Spatial gene expression maps for Yap, Cyr61, Foxa2 and Ptch1. (Bottom) 1D gene expression profiles along the tissue compartments boundary for the same transcripts. **C:** (Top) Spatial gene expression maps for Nkx2.2, Nkx6.1, Pax7 and Pax3. (Bottom) 1D gene expression profiles along the tissue compartments boundary for the same transcripts.

